# Cellular visualization of G-quadruplex RNA via fluorescence lifetime imaging microscopy

**DOI:** 10.1101/2023.05.07.539224

**Authors:** Jenna Robinson, Stine G. Stenspil, Karolina Maleckaite, Molly Bartlett, Marco Di Antonio, Ramon Vilar, Marina K. Kuimova

## Abstract

Over the last decade, appreciation of the roles of G-quadruplex (G4) structures in cellular regulation and maintenance have rapidly grown, making the establishment of robust methods to visualize G4s increasingly important. Fluorescent probes are commonly used for G4 detection *in vitro*, however, achieving sufficient selectivity to detect G4s in a dense and structurally diverse cellular environment is challenging. The use of fluorescence probes for G4 detection is further complicated by variations of probe uptake into cells, which may affect fluorescence intensity independently of G4 abundance. In this work, we report an alternative small-molecule approach to visualize G4s that does not rely on fluorescence intensity switch-on and thus, does not require the use of molecules with exclusive G4 binding selectivity. Specifically, we have developed a novel thiazole orange derivative, TOR-G4, that exhibits a unique fluorescence lifetime when bound to G4s compared to other structures, allowing G4 binding to be sensitively distinguished from non-G4 binding, independently of local probe concentration. Furthermore, TOR-G4 primarily co-localizes with RNA in the cytoplasm and nucleoli of cells, making it the first lifetime-based probe validated for exploring the emerging roles of RNA G4s *in cellulo*.

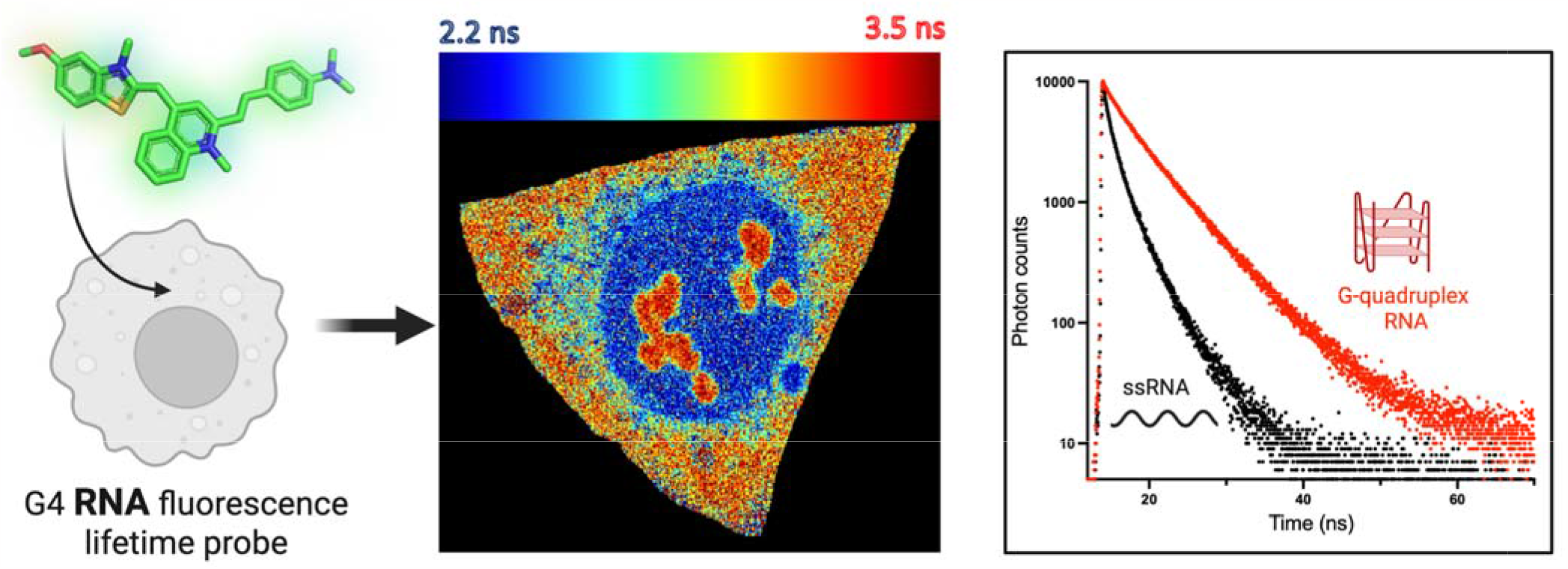

## INTRODUCTION

The structural landscape of DNA and RNA is essential in facilitating the cellular function of nucleic acids, making nucleic acid secondary structures critical for regulation of many cellular processes.^1–3^ G-quadruplexes (G4s) are secondary structures that form in DNA and RNA through non-canonical hydrogen bonding between four guanine bases.^4,5^ As understanding of pathways regulated by nucleic acids has evolved, G4s have progressed from being considered as elusive structures that form in repeat sections of telomeres,^6^ to prospective regulators of cell biology that form extensively across cells.^4,5^

In the nucleus, DNA G4s have been associated with epigenetic regulation of gene expression, through their interactions with regulatory proteins, such as transcription factors and chromatin modifiers.^7,8^ Whilst RNA G4s have been linked to regulation of RNA splicing, transport and translation, as well as RNA-mediated stress responses in the cytoplasm.^9–11^ As many of these processes occur in spatially confined regions of the cell,^9–11^ deciphering the biological roles of G4s now requires visualization not just within individual cells, but within restricted subcellular compartments. It is thus necessary that the tools we use to interrogate G4 formation continue to evolve as our study of G4 biology becomes simultaneously more wide-ranging and precise.

The first direct proof of G4 formation within cells was provided using G4-specific antibodies visualized by immunostaining.^12–14^ In parallel, several small-molecule ‘switch-on’ probes have been reported that become fluorescent upon binding to G4s.^15–17^ Whilst such fluorescence intensity probes are very useful for *in vitro* studies, their use for understanding G4 biology in cells presents several challenges. Exceptionally high G4 selectivity is required for such probes to work effectively within cells, due to the large abundance of non-G4 secondary structures that may result in a smaller ‘switch-on’ which is, however, sufficient to produce false positive fluorescent signals.^2,3^ Not only is this high G4 binding affinity difficult to achieve, but may in fact alter natural G4 formation within cells.^18^ Secondly, fluorescence intensity is intrinsically concentration dependent,^19^ and so may vary independently of G4 content due to differences of probe uptake into a given cell or organelle.

To address the limitations of fluorescence intensity probes, alternative methods have arisen to visualize G4s based on changes to the fluorescence lifetime of a molecule upon nucleic acid binding.^20–23^ For environmentally sensitive fluorophores, the rate of fluorescence decay can vary depending on the binding conformation and the environment of a molecule.^24^ Such variations in fluorescence lifetime make it possible to identify binding to specific structures, even when using less selective molecules that interact promiscuously with multiple topologies (Figure 1A). Additionally, fluorescence lifetime is generally concentration-independent and therefore remains constant regardless of cellular uptake (Figure 1A).^24^ Currently, there are a limited number of fluorescence lifetime-based probes that have been reported for visualizing G4s,^20–23^ which have enabled understandings of G4 dynamics within cells such as G4 unfolding by the action of DNA helicases.^21^ However, previous G4 lifetime probes were developed for DNA G4s, which has left the emerging role of RNA G4s inaccessible for study using lifetime-based approaches.

**Figure 1.**
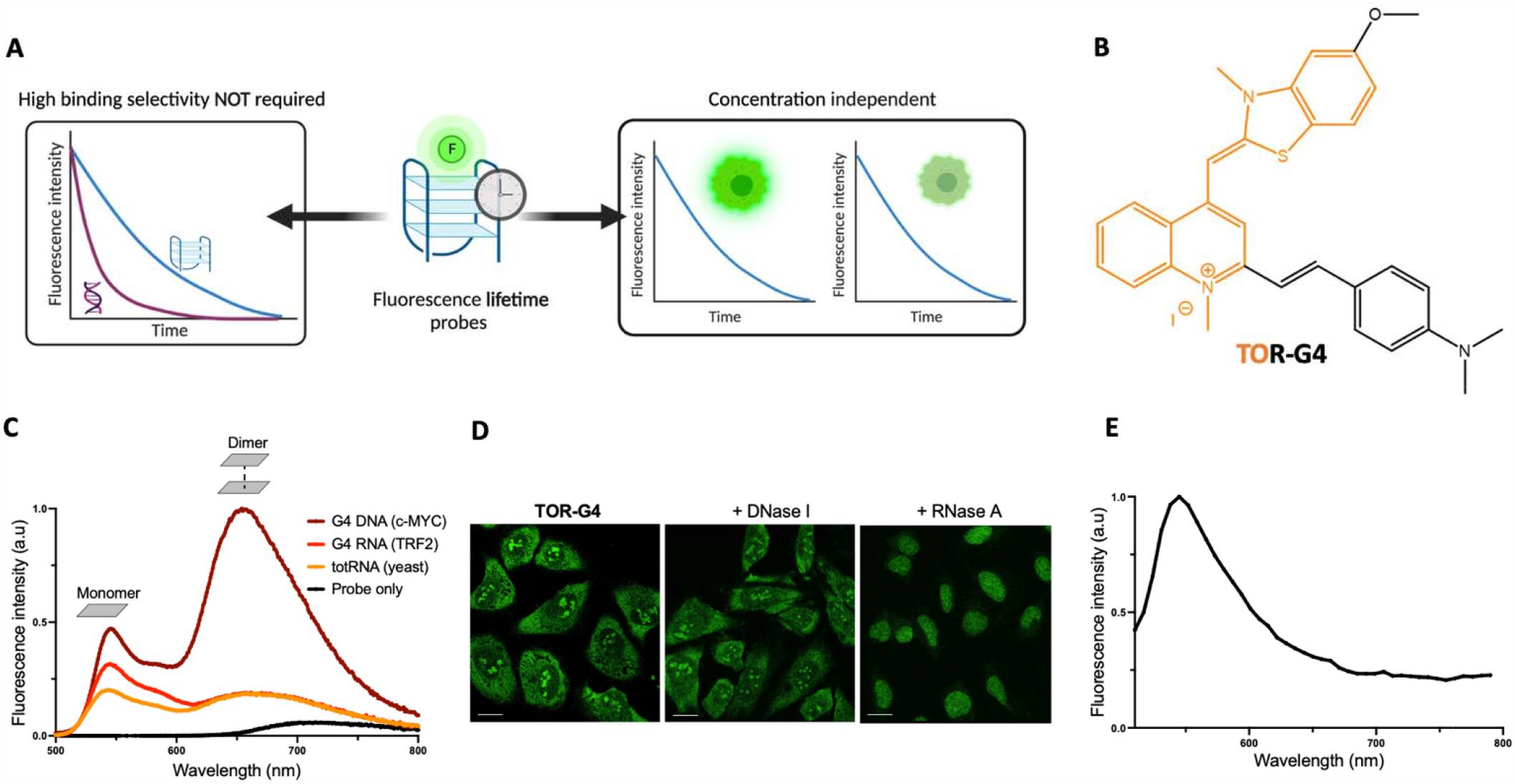
A) Schematic of the principles and benefits of using fluorescence lifetime approaches for G4 imaging. Binding t each nucleic acid topology is characterized by a unique time-resolved decay and lifetime e.g. longer for G4 (blue), shorter for duplex (purple). Therefore, each topology can be clearly detected even in the presence of the other, and high binding selectivity is not required. Additionally, the measurement is independent of probe concentration. B) Structure of thiazole orange derivative **TOR-G4** - thiazole orange core is shown in orange. C) *In vitro* emission spectra of **TOR-G4**, free in solution (black) and bound to G4 DNA (maroon), G4 RNA (red) and total RNA (orange), following 470 nm excitation. D) Confocal images of **TOR-G4** in fixed U2OS cells before and after treatment with DNase I or RNase A. The RNase treatment causes significant changes in the probe’s distribution in cells. Scale bar = 20 μm. E) Cellular emission spectrum of **TOR-G4** in U2OS cells following 477 nm excitation.

In this work, we designed TOR-G4 - a derivative of thiazole orange (Figure 1B), and explored its use as a new G4 fluorescence lifetime probe. Thiazole orange (TO) was selected as the core motif of the probe due to its reported selectivity for G4s.^25–28^ Additionally, TO is known to be an environmentally sensitive probe where rotation around the bridging single bond leads to fast relaxation of the excited state and thus, a short fluorescence lifetime; ^29,30^ however, on binding to DNA, this rotational freedom is limited resulting in a significant increase in fluorescence intensity and lifetime.^29,30^ The lifetime sensitivities of TO thus make it a promising starting point to develop new lifetime probes for G4 imaging.

Here, we demonstrate that the lifetime of **TOR-G4** is highly dependent on the interacting nucleic acid structure – being highest in the presence of G4s and lower for other sequences. Within cells, we showed that the probe primarily co-localizes with RNA and displays fluorescence lifetimes which are dependent on G4 binding, making it the first lifetime probe suitable for detecting RNA G4s within cells. Overall, we present **TOR-G4** as a novel molecule for visualizing G4 RNA that overcomes many of the limitations of intensity-based switch-on probes.

## RESULTS

### Characterization of a new thiazole orange derivative

Several previous reports have shown that structural modifications of TO can lead to increased G4 selectivity.^25,27,28^ Thus, we designed and synthesized **TOR-G4** (Figure 1B) which includes the addition of a benzylstyryl unit to extend electron conjugation of the original TO motif and thus, its potential for ***π*-**stacking with G-quartets. ^25,28^ We also considered that an ideal G4 probe could be used for imaging within both cells and tissues, the latter of which requires two-photon excitation (TPE) of samples. In TPE, the combined energy of two photons is used to excite a sample, which allows for the use of longer wavelengths of light that scatter less in tissue. The presence of an electron donor-acceptor-donor structure has previously been shown to enhance the two-photon absorption propensity of molecules.^31,32^We therefore introduced a methoxy and dimethylamine group to either end of the probe (chosen due to commercial availability of starting materials), to sandwich the positively charged nitrogen in the center of the molecule and in turn optimize the probe’s suitability for TPE.

After successful synthesis and characterization of **TOR-G4** (Figures S1-7), we next investigated the photo physics of the molecule in the absence and presence of various nucleic acid structures (Figures 1C, S8, Table S1). In aqueous media, the probe displays a very weak emission centered at ca. 700 nm, however, the addition of nucleic acids results in a significant increase in the fluorescence intensity with two spectral peaks visible, centered at 540 nm and 660 nm (Figure 1C). As TO is known to self-assemble into aggregates in aqueous solution,^33–37^ we hypothesized that the two emission peaks were due to the monomer and dimer/aggregated form of the probe.

To investigate probe aggregation, we first measured the excitation spectra at both peaks and observed distinct spectra corresponding to separate ground state entities (Figure S9). The species emitting at 660 nm had a blue-shifted excitation maximum relative to the species emitting at 540 nm, which is characteristic of H-aggregate formation.^38^ To further confirm that the 660 nm peak arises from probe aggregates, we performed a concentration titration where the concentration of TOR-G4 was increased in the presence of the DNA G4 sequence *c-MYC* (Figure S9). We observed that the intensity of the 660 nm peak increased rapidly with increasing probe concentration. This change in peak ratios shows that the species emitting at 660 nm is promoted by higher concentrations of TOR-G4 and is therefore likely to correspond to the aggregated form of the probe (which also aligns with previous work on TO aggregation).^37^ The monomer form of the probe (emitting at 540 nm) only appears after the addition of oligonucleotides, potentially due to stronger interactions of the probe with DNA/RNA than with itself (Figures 1C).

### TOR-G4 localizes with RNA within cells

We next set out to characterize the cellular uptake and localization of the molecule. We opted to characterize **TOR-G4** within fixed cells, which are commonly used in the field when imaging G4s via immunofluorescence,^12,13,39^ due to the observed toxicity of the molecule (Figure S10, GI_50_ = 143 nM). Confocal microscopy images of the probe within U2OS cells revealed a distinct staining pattern, with particularly high intensity in the cytoplasm and nucleoli (Figure 1D). We hypothesized that this staining pattern may be indicative of RNA binding, as nucleoli are the site of ribosomal RNA transcription and are also highly guanine-rich.^40^ This notion was supported by the analogous staining of **TOR-G4** to a commercial RNA stain (SYTO RNASelect, Figure S11). To confirm the preferential binding of **TOR-G4** to RNA, we performed nuclease experiments where cells were treated with: i) DNase I which removes cellular DNA; ii) RNase H which degrades DNA:RNA hybrids; iii) RNase A which degrades single stranded RNA and DNA:RNA hybrids; iv) RNase T1 which cleaves single stranded RNA only. We found that treatment with DNase I and RNase H did not substantially alter the localization nor the fluorescence intensity of the probe (Figure 1D, Figure S12). In contrast, treatment with RNase A and T1 resulted in a significant drop in the fluorescence intensity and a change in the staining pattern of the probe, leaving solely nuclear staining without characteristic nucleolar staining (Figure 1D, S12). In addition to this, transcriptional inhibition of cells via addition of the RNA polymerase inhibitor DRB resulted in a large reduction of nucleolar staining (Figure S13) – demonstrating the high intensity of the probe in nucleoli is indeed due to active transcription in this region.

The cellular localization results suggest that **TOR-G4** naturally localizes with RNA in the cytoplasm and nucleoli of cells and is not significantly affected by the removal of DNA or DNA:RNA hybrids. However, in the absence of RNA, the probe binds to DNA, in turn becoming a nuclear stain. This preferential colocalization of **TOR-G4** with RNA makes the molecule one of a limited number of probes well suited for studying G4 RNA within cells using fluorescence microscopy.^41–44^

Interestingly, the fluorescence spectrum of **TOR-G4** within U2OS cells revealed only the presence of the monomer species emitting at 540 nm (Figure 1E). The exclusive existence of the monomer species in cells may be due to the high density of interacting nucleic acids. The high cellular ion concentration may also contribute to probe disaggregation, as we found high potassium concentration reduced dimer emission, potentially via G4 stabilization. (Figure S14).

We considered that probe disaggregation may also occur upon binding to other biomolecules that are present in cells, such as proteins and lipids. To investigate interactions with other cellular components, we measured the emission spectrum of the probe in U2OS cell lysate before and after the treatment with nuclease. We found that in total lysate, two emission peaks at ∼550 nm and 650 nm were observed in the spectrum (Figure S15), consistent with the presence of both the monomer and the aggregate species. In contrast, after treatment with nuclease, only a single peak corresponding to the probe aggregates (emitting at ∼650 nm) was observed. This demonstrates that to observe the monomer species, specific binding to nucleic acids is required. The probe is thus well suited for specifically studying nucleic acids within cells without interference from other biomolecules.

### Fluorescence lifetime of TOR-G4 varies based on nucleic acid structure

Having confirmed that **TOR-G4** stains RNA in cells and emits from its monomer state upon binding, we set out to test whether it may be used as a fluorescence lifetime probe for G4s. To this end, we measured the time-resolved fluorescence traces of the cellularly-relevant monomer species in the presence of multiple structures, including six G4-forming sequences, considering both RNA and DNA structures. As a baseline measurement, we also measured the probe’s lifetime in the presence of commercially available total RNA (totRNA) extracted from yeast, which contains a mixture of all RNA structures present in a typical cell. Similarly, we tested yeast tRNA cell extract as an additional subset of mixed RNA topologies which may include G4s.^45^ Total RNA extracted from human U2OS cells was also tested to ensure the structural diversity of RNA in yeast adequately reflects that found within human cells. Next, we selected RNA sequences that cannot form G4s, such as simple single stranded RNA sequences which lack secondary structure, as well as a hairpin RNA sequence and stem-loop structures with symmetric (cov-bulge) or asymmetric (ucu-bulge) bulges. Finally, we recorded the lifetime of **TOR-G4** bound to duplex DNA extracted from calf thymus to consider lifetime selectivity for RNA over DNA.

We found that the time-resolved fluorescence decays of **TOR-G4** fit well to a bi-exponential decay function (Figures 2A, S16, S17, Table S3). Both lifetime components were highly dependent on the nucleic acid structure the probe interacted with, being significantly higher in the presence of G4 structures than for mixed or non-G4 topologies. In the presence of G4s, the decay of **TOR-G4** also had higher contributions from the longer lifetime component (amplitude 2 and tau 2, Figure S17), causing the probe to have a significantly higher intensity-weighted average lifetime when bound to G4s (4-6 ns) compared to non-G4/mixed topologies (2-3 ns) (Figures 2A, B). The fluorescence enhancement of the probe upon DNA and RNA binding also displayed similar trends to that of the fluorescence lifetime, being the highest when the probe was bound to G4s – although there was significantly higher variance in the intensity-based measurements (Figure S18).

**Figure 2.**
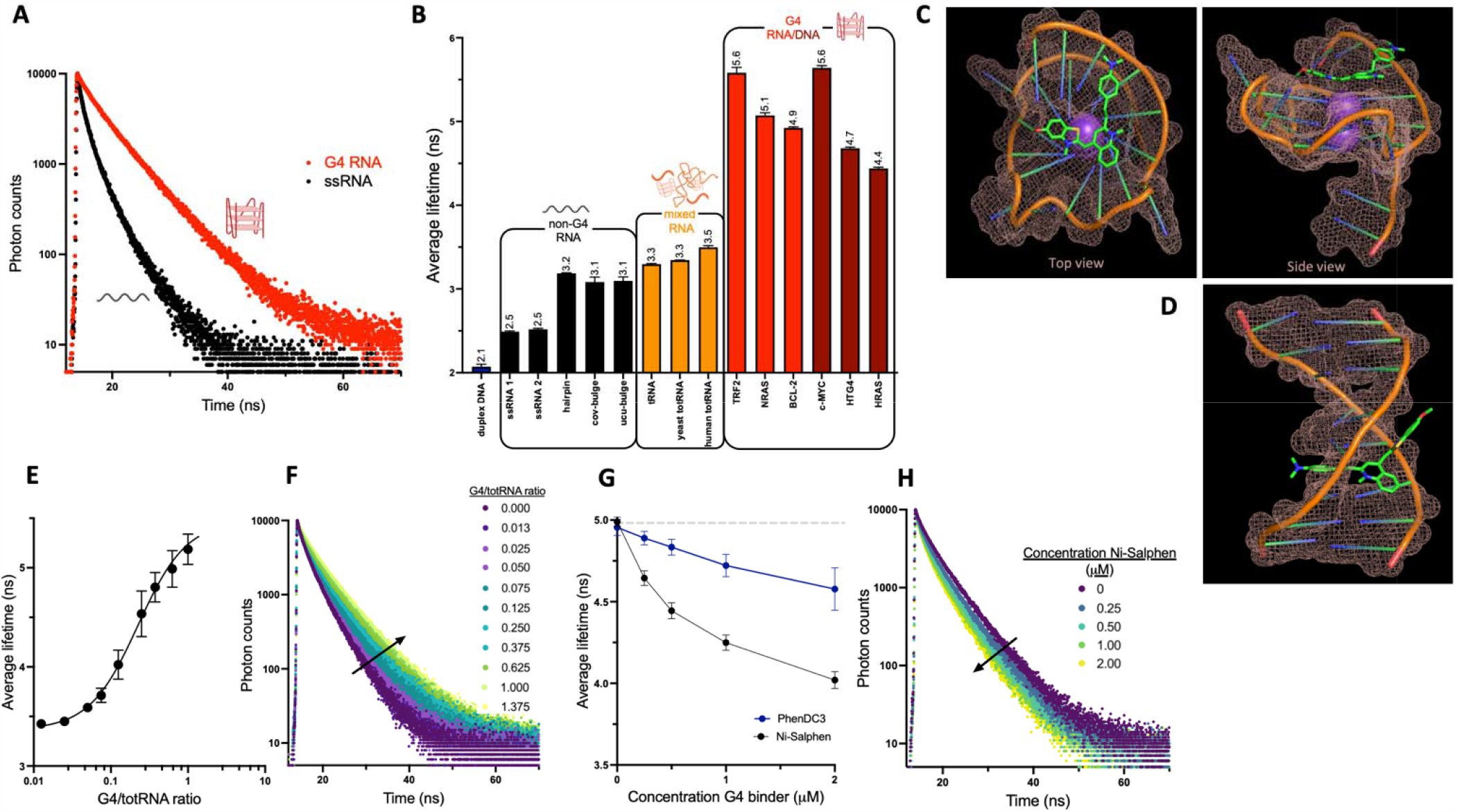
A) Sample time-resolved fluorescence decays of **TOR-G4** when bound to a G4 or a single-stranded RNA sequence. B) Average fluorescence lifetimes of **TOR-G4** when bound to various nucleic acid structures after excitation at 467 nm and detection at 540±32 nm. C) Molecular modelling of **TOR-G4** bound to the G-quadruplex *c-MYC* or D) a duplex DNA structure, demonstrating variations in the probe’s binding conformation. E) Average fluorescence lifetimes and F) time-resolved fluorescence decays of **TOR-G4** recorded in the presence of varying ratios of G4 RNA to totRNA. G) Average fluorescence lifetimes recorded from displacement assays – displacing **TOR-G4** from an RNA G4 sequence with established G4 binders Ni-Salphen and PhenDC3 (see Figure 4 for molecular structures). H) Time-resolved fluorescence decays of **TOR-G4** during G4 displacement assay with Ni-Salphen. Error bars are standard deviation for experiments performed in tripl cate.

Despite preferential cellular binding to RNA, the lifetime values for G4 DNA and RNA were comparable, suggesting that **TOR-G4** has selectively high lifetimes for all G4 structures regardless of the nucleic acid. To further test if the lifetime of **TOR-G4** was dependent on the formation of G4s, lifetime measurements were made with the parallel G4 sequence c-MYC, in binding buffer containing LiCl (rather than KCl), which does not significantly stabilize G4s.^46^ Destabilization of G4s by removal of K^+^ ions resulted in a significant drop in the measured fluorescence lifetime of the probe by 1.3 ns (Figure S19), showing G4 formation is essential for obtaining a high probe lifetime. We noted that the lifetime in lithium-containing buffer was still higher than that of non-G4 sequences, suggesting some G4 formation was still occurring even in the absence of added K^+^ ions. To confirm this, CD spectra of c-MYC in both K^+^ and Li^+^ buffer were recorded, which revealed the parallel G4 signature (negative and positive peaks at −240 and +265 nm respectively, Figure S19)^47^ were present in both condi**t**ions, thus explaining the maintenance of a relatively high lifetime in Li^+^ buffer.

To rationalize the structural dependence of **TOR-G4**’s lifetime, we performed molecular modelling studies comparing the DFT-optimized conformation of the probe when bound to the structure that produced the lowest lifetime (duplex DNA) and the highest lifetime (the G4 *c-MYC*) (Figures 2C, D). Docking studies showed that when interacting with the duplex structure, a portion of the probe intercalates in between base pairs, however, due to the extended size of the molecule, part of the probe is forced to point away from the DNA backbone. This displaced fragment likely has a substantial amount of rotational freedom, which is likely to increase the rate of non-radiative decay and result in a lower probe lifetime.^24^ In contrast, when bound to the G4 structure via end-stacking, the entirety of the probe fits on top of the G-tetrad. Such a binding arrangement means that the whole of the molecule is involved in the -stacking interaction with the G4 and is thus more conformationally restricted, likely leading to a longer lifetime.

To explore the hypothesis that differences in lifetime are explained by distinct binding interactions, we performed binding affinity titrations on selected sequences (Figures S20 and S21). Both the monomer and dimer peak were found to change with increasing nucleic acid concentration, suggesting both forms of the probe may interact with DNA and RNA. For the monomer species, we found the **K**_a_ of **TOR-G4** was approximately 3× higher for G4 sequences than for other structures. Interestingly, we also noted that the binding affinity for duplex DNA was comparable to that of totRNA – suggesting that the cellular localization with RNA is not due to specific RNA binding preferences. Instead, within cells intercalation into DNA may be impaired due to chromatin structure, as has been previously described for many intercalating dyes,^48–50^ causing the probe to interact primarily with RNA which is abundant in the nucleoli and cytoplasm.

### Perturbing G4 binding alters lifetime of TOR-G4

Having established that the lifetime of **TOR-G4** varies depending on nucleic acid structure, we next tested if the lifetime of the probe could dynamically respond to changes in G4 binding. We opted for two perturbation strategies to either increase or decrease G4 binding. Firstly, to assess the lifetime response to increasing G4 prevalence, we measured the time-resolved decays of **TOR-G4** in a solution of yeast totRNA and investigated how the addition of an RN**A** G4 sequence affected the detected lifetime of the molecule. We found that the lifetime gradually increased from 3.5 to 5.5 ns with increasing G4 concentration (Figures 2E, F), demonstrating the probe’s ability to detect elevated G4 prevalence even in the presence of many other RNA structures.

Secondly, to investigate how the probe responds to reduced G4 binding, we performed a G4 displacement assay using an RNA G4 sequence. To achieve this, we used two validated G4 ligands that are structurally distinct, PhenDC3 ^51,52^ and Ni-Salphen^53^ (Figures 4A,B) and titrated them into a solution of **TOR-G4** pre-bound to an RNA G4. As expected, we observed that increasing the concentration of G4 binders displaced the probe from its RNA G4 substrate, resulting in a significant drop in fluorescence lifetime, by 1 ns with Ni-Salphen and 0.4 ns with PhenDC3 (Figure 2G,H). The greater lifetime reduction obtained with Ni-Salphen also aligns with the reported higher binding of Ni-Salphen to G4s compared to PhenDC3.^52,54^ In contrast, when the same experiment was repeated using a hairpin RNA sequence as the **TOR-G4** binding substrate, we did not observe any significant change in the probe lifetime (Figure S22). Additionally, displacement from yeast totRNA resulted in a modest drop in lifetime (by 0.3 and 0.1 ns when adding Ni-Salphen and PhenDC3, respectively, Figure S22), which is in agreement with total RNA containing a mixture of G4 and non-G4 structures. Overall, these experiments demonstrate that the lifetime of **TOR-G4** is highly sensitive to G4 binding and may be used to assess G4-prevalence within cells.

### TOR-G4 detects G4 content in cells via FLIM

After the *in vitro* validation of **TOR-G4** as a successful G4 fluorescence-lifetime probe, we next considered its suitability for detecting G4 abundance in a cellular environment via fluorescence lifetime imaging microscopy (FLIM). Here, the intensity-weighted average fluorescence lifetime is calculated for individual pixels of a confocal image to yield a colour-coded map of pixel lifetimes, where high lifetime pixels are shown in red and lower lifetime pixels in blue (Figure 3A). FLIM analysis of **TOR-G4** in U2OS cells, revealed that the lifetime of the probe varies considerably across the cell (Figure 3A-C, Figure S23).

**Figure 3.**
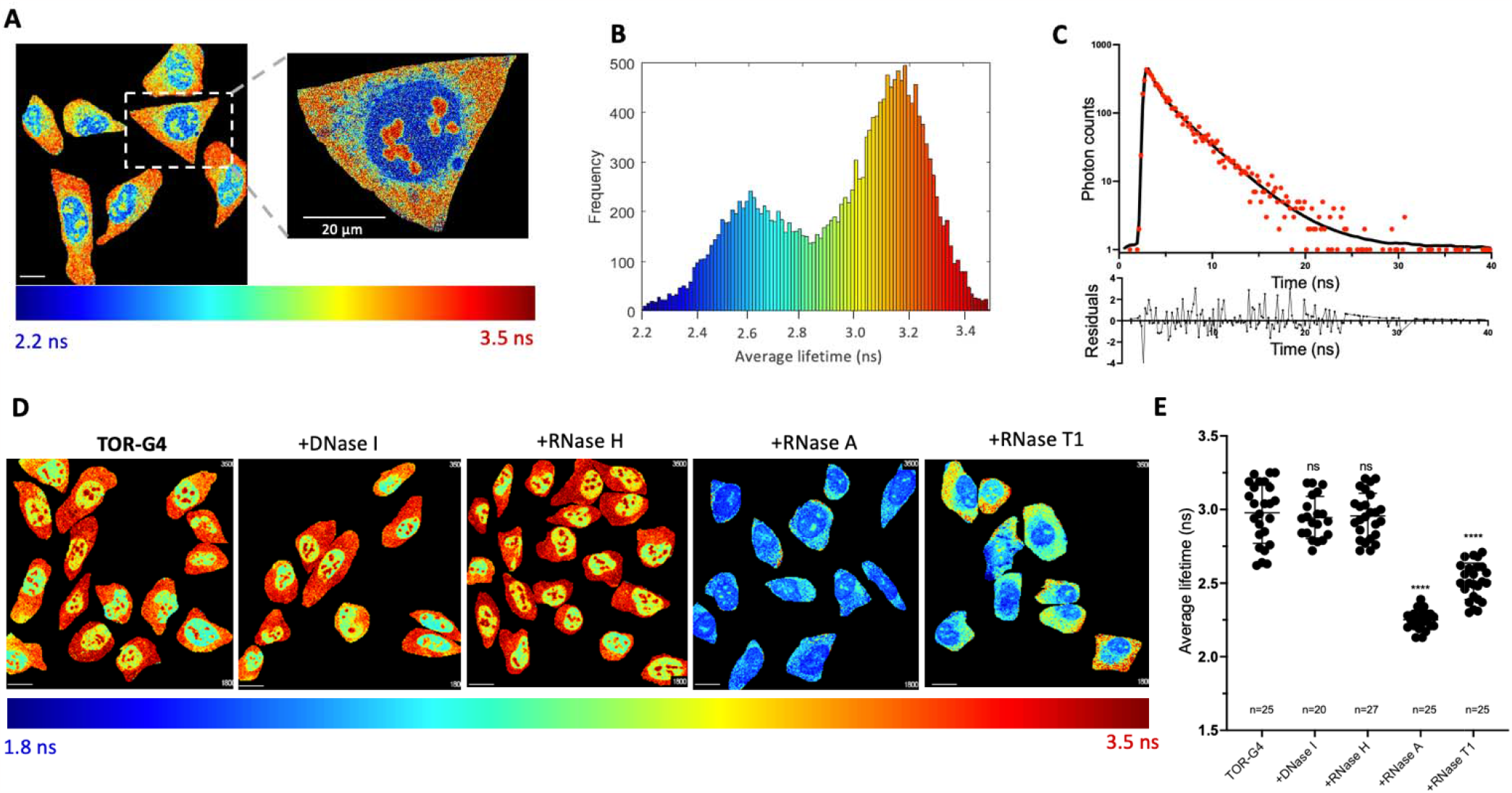
A) FLIM images of **TOR-G4** in U2OS cells. Overview image was taken at 256×256 pixel resolution and zoomed image of cell was separately acquired and segmented at 512×512 pixel resolution. B) Histogram of pixel frequency against average fluorescence lifetime of **TOR-G4** within U2OS cells. C) Example of cellular fluorescence decay and decay fit residuals of **TOR-G4** to a biexponential decay function. Decay taken from a pixel in the nucleoli. D) FLIM images and E) average fluorescence lifetime of **TOR-G4** in cells following treatment with nucleases. Excitation at 477 nm and emission collected across 550-700 nm. Scale bars = 20 m.

Specifically, there were two distinct lifetime environments, identified both by fitting lifetime decays (Figure 3A,C) and by phasor analysis of images (Figure S24). The first probe environment was at ~3.2 ns in the cytoplasm and nucleoli, which is consistent with the *in vitro* measurements of the probe bound to totRNA (i.e. representing a mixture of G4 and non-G4 interactions). In comparison, a significantly lower lifetime is seen for the non-nucleolar parts of the nucleus, where there is a peak lifetime of 2.6 ns. This lower lifetime is comparable to that measured with single stranded RNA *in vitro* and may be due to binding of the probe to mRNA transcripts in the nucleus. We also noted similar lifetime distributions were obtained when imaging the **p**robe via two-photon excitation (Figure S25), thus demonstrating **TOR-G4** may be suitable for tissue imaging in the future.

To further validate the lifetime measured within cells specifically arises from probe binding to RNA, the lifetime of **TOR-G4** was measured before and after treatment with DNase I as well as RNase H, A and T1 (Figure 3D,E). Similarly, to the fluorescence intensity measurements, the fluorescence lifetime of **TOR-G4** was only found to be significantly diminished after treatment with RNase **A** and RNase T1. Inhibition of transcription via treatment with DRB also resulted in a reduction in fluorescence lifetime (F**i**gure S13), thus further indicating that the fluorescence lifetime of the probe primarily arises from its interaction with RNA in cells.

We next sought to validate that the lifetime measured within each whole cell does not vary based on the probe uptake. To assess this, we measured the average fluorescence intensity within each cell at two probe concentrations (2 M and 5 M) and correlated it with the average cellular lifetime. Whilst changing probe concentration results in large changes in the fluorescence intensity measured within cells, we found no correlation between the average intensity of a cell and its fluorescence lifetime (Figure S26). We additionally found that the fluorescence lifetime of the probe remained consistent (< 0.1 ns change) after irradiation of light and continuous imaging across 6 hours (Figure S27). These results demonstrate that the fluorescence lifetime of **TOR-G4** is robust and not significantly affected by fluctuations in cellular probe uptake or light exposure, unlike fluorescence intensity measurements.

Finally, we tested the sensitivity of the probe towards G4 binding by conducting two perturbation experiments in cells. Firstly, we performed cellular G4 displacement assays where, similarly to the *in vitro* assay, two G4 binders (PhenDC3 and Ni-Salphen, 1 μM, Figure 4A-D) were added to cells which had been pre-incubated with **TOR-G4**. After 4 hours of incubation with the corresponding G4 binders, significant drops in the fluorescence lifetime were observed (F**i**gure 4C,D), with Ni-Salphen resulting in a larger drop in lifetime compared to PhenDC3 – thus showing the same trend as the *in vitro* displacement results.

**Figure 4.**
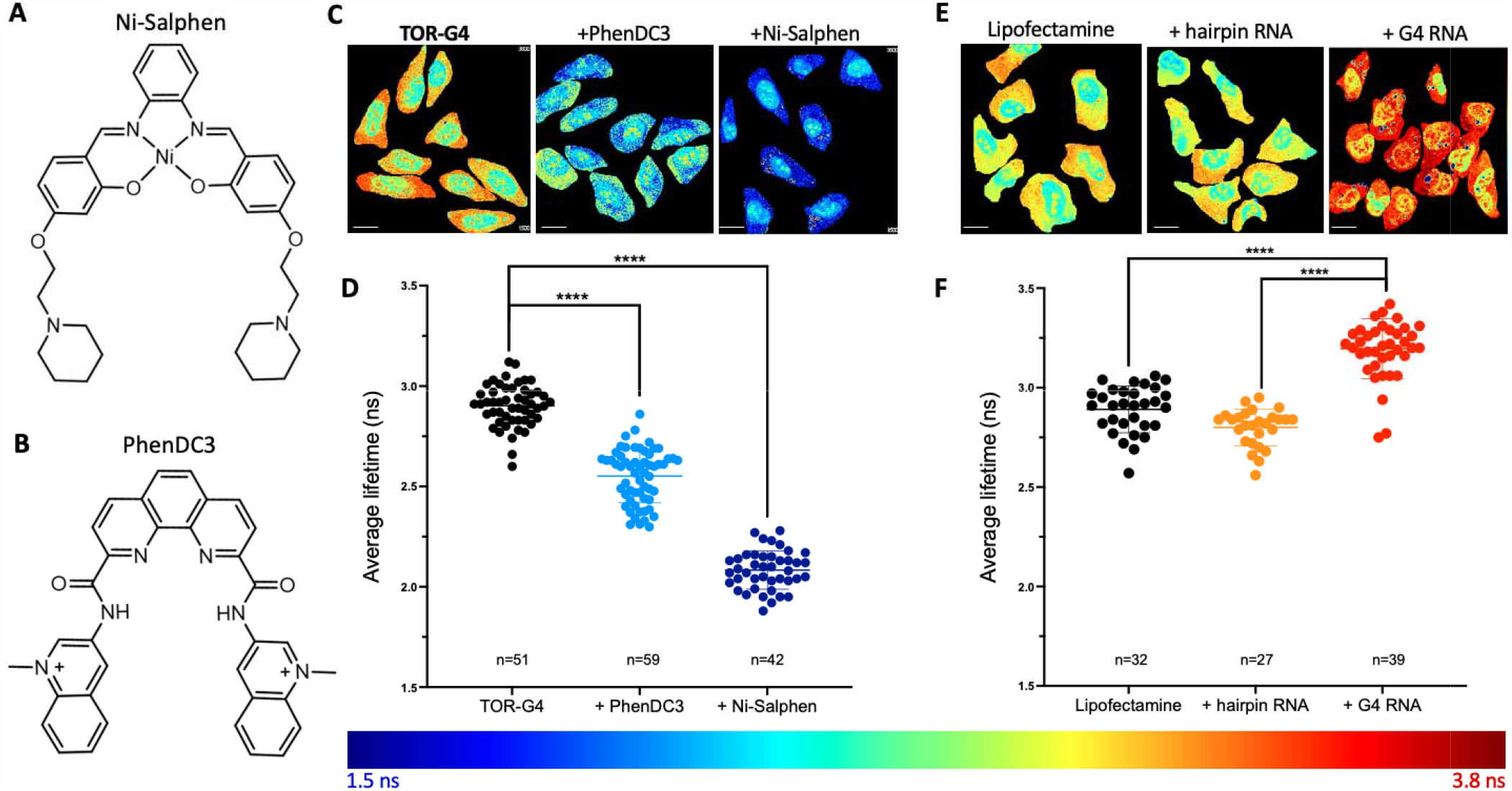
A) Structure of Ni-Salphen and B) PhenDC3. C) FLIM images and D) average fluorescence lifetime of **TOR-G4** in U2OS cells after displacement with G4 ligands Ni-Salphen and PhenDC3. E) FLIM images and F) average fluorescence lifetime of **TOR-G4** in U2OS cells transfected with lipofectamine only, hairpin RNA or G4 RNA. Excitation at 477 nm and emission collected across 550-700 nm. Scale bars = 20 m.

Secondly, we investigated the probe’s response to increased G4 abundance by transfecting G4 RNA into cells. As a control we also measured the lifetime of **TOR-G4** within cells that had only been treated with the transfecting agent (lipofectamine 2000) or alternatively, transfected with a hairpin RNA sequence. As expected, transfection with the G4-forming sequence resulted in a significantly higher measured lifetime compared to lipofectamine treatment alone or hairpin RNA transfection (Figure 4E,F). These results demonstrate that both decreasing and increasing G4 binding within cells results in significant changes in the lifetime of **TOR-G4**, making the probe an effective and concentration-independent tool for monitoring changes in cellular G4 formation.

## DISCUSSION

The formation of G4 structures in DNA and RNA has been linked to multiple regulatory pathways across the cell, in increasingly diverse and compelling ways.^7–11^ G4s are thus promising therapeutic targets for diseases including cancer and neurodegeneration.^55,56^ Untangling the role of G4s in cellular processes has been facilitated by G4 visualization using either small-molecule fluorescent probes or fluorescently-labelled antibodies.^12–17,39,57^

Historically, research on G4s has been focused on DNA G4s which is reflected in the greater availability of fluorescent probes that exhibit nuclear staining.^58^ In comparison, visualization of RNA G4s (rG4s) is still a relatively new area of study.^41–44^ Previously, it was suggested that RNA G4s may even be globally unfolded within cells due to the persistent activity of G4 helicases.^59^ However, more recent studies have provided extensive evidence that rG4s not only exist within cells but may be pivotal for mediating RNA interactions with proteins that control processes such as splicing, translation and stress granule formation.^9–11^

Despite the exciting emerging roles of RNA G4s within cells, imaging rG4s poses a unique challenge due to the increased structural diversity of RNA compared to DNA.^1^ This means that any rG4 probes must exhibit exceptionally high binding selectivity for G4s. Additionally, as rG4s are often found in phase-separated regions of the cell (and may even drive phase separation events),^60–62^ probe uptake must be considered when interpreting fluorescence images. For example, differences of probe uptake into phase-separated organelles confound measurements of fluorescence intensity, where increased fluorescent signal may be due to local probe concentration rather than G4 abundance.

Fluorescence lifetime imaging microscopy can be used to overcome stringent requirements for G4 binding selectivity, as it allows for binding interactions with G4s to be distinguished from that of other structures. Furthermore, lifetime measurements provide a readout of G4 abundance which is independent of local probe concentration. Whilst some probes have been reported to display distinct fluorescence lifetimes when bound to G4 DNA compared to other structures,^20–23^ lifetime probes for RNA G4s have yet to be reported. In this work, we have filled this gap by describing a fluorescence lifetime-based probe (**TOR-G4**) suitable for imaging RNA G4s in cells.

We first characterized the fluorescent properties of **TOR-G4**, noting that *in vitro* the probe can exist in either a monomeric or an aggregated form, whilst in cells the molecule fully disaggregates, presumably upon nucleic acid binding. The fluorescence lifetime of the monomer species is highly dependent on the structure of the interacting nucleic acids, being the highest when bound to G4 structures, compared to non-G4 or mixed topologies. These differences in the probe’s lifetime were rationalized with molecular modelling studies showing that the conformational flexibility of **TOR-G4** is restricted when bound to a G4, compared to a duplex structure, which would in turn reduce the rate of non-radiative decay.

Unlike other G4 lifetime probes, within cells **TOR-G4** predominantly stains the cytoplasm and nucleoli and is sensitive to treatment with RNase. Despite this, *in vitro* testing revealed there was no binding selectivity for RNA over DNA. Instead, we hypothesize that the cellular RNA selectivity of **TOR-G4** is due to its intercalating binding mode. Previous work has shown that many small molecules that intercalate into DNA *in vitro*, are not able to achieve intercalation when DNA is condensed into chromatin, leading solely to RNA binding.^48–50^ This is in contrast to other DNA G4 probes that have been shown to interact with DNA via groove binding,^23^ which allows for DNA binding even in chromatin. The RNA selectivity of **TOR-G4** within cells in turn makes the probe best suited for studying RNA G4s.

To validate the sensitivity of **TOR-G4** to G4 content in cells, we performed G4 displacement assays with known G4 ligands PhenDC3 and Ni-Salphen and also measured the lifetime of the probe after G4 RNA transfection. We demonstrated that displacing **TOR-G4** from G4s results in a large and significant drop in the average cellular fluorescence lifetime. Similarly, inducing G4 formation via RNA transfection resulted in a significant increase in probe lifetime compared to transfection of non-G4 structures. Together these results confirm that the molecule can be used as a sensitive tool to probe cellular rG4 abundance.

## CONCLUSION

In this work we present the characterization and application of **TOR-G4** – a small-molecule suitable for imaging RNA G4s via FLIM. By characterizing the fluorescence lifetime of **TOR-G4** with a range of G4 and non-G4 structures, we established that the probe displays distinct fluorescence lifetimes when bound to G4s compared to other structures, which can be perturbed by the addition of known G4 ligands. In cells, the probe primarily localizes in cellular compartments that are rich in RNA, making it one of a limited number of molecules suitable for probing RNA G4 formation. Overall, **TOR-G4** represents an alternative, concentration-independent tool for visualizing RNA G4s, which overcomes the need to obtain exclusive G4 binding selectivity within cells.

## ASSOCIATED CONTENT

### Supporting Information

Description of experimental protocols, materials and supplementary figures (S1-S27) as mentioned in the text (PDF).

## AUTHOR INFORMATION

### Author Contributions

J.R., M.K.K. and R.V. designed the study with input from M.D.A. on cellular experiments. J.R., M.D.A., M.K.K., and R.V. co-wrote the paper. J.R., S.G.S., K.M. and M.B. performed experiments and analyzed the data. M.D.A., M.K.K. and R.V. secured the funding for this work.

## ACKNOWLEDGMENTS

J.R. is funded by the Engineering and Physical Sciences Research council (EPSRC, grant number EP/S023518/1) and the NIHR imperial biomedical research centre. M.B. is funded by the Engineering and Physical Sciences Research council (EPSRC, grant number EP/S023232/1). M.D.A is supported by a Biotechnology and Biological Sciences Research Council (BBSRC) David Phillips Fellowship [BB/R011605/1] and is a recipient of the Lister Institute Research Prize.

## ABBREVIATIONS

G4: G-quadruplex rG4
RNA: G-quadruplex
K_a_: Association constant
FLIM: Fluorescence Lifetime Imaging Microscopy

## REFERENCES

(1) Ganser, L. R.; Kelly, M. L.; Herschlag, D.; Al-Hashimi, H. M. The Roles of Structural Dynamics in the Cellular Functions of RNAs. Nat. Rev. Mol. Cell Biol. 2019, 20 (8), 474–489. 10.1038/s41580-019-0136-0.

(2) Kaushik, M.; Kaushik, S.; Roy, K.; Singh, A.; Mahendru, S.; Kumar, M.; Chaudhary, S.; Ahmed, S.; Kukreti, S. A Bouquet of DNA Structures: Emerging Diversity. Biochem. Biophys. Reports 2016, 5, 388–395. 10.1016/J.BBREP.2016.01.013.

(3) Morris, K. V.; Mattick, J. S. The Rise of Regulatory RNA. Nat. Rev. Genet. 2014, 15 (6), 423–437. 10.1038/nrg3722.

(4) Spiegel, J.; Adhikari, S.; Balasubramanian, S. The Structure and Function of DNA G-Quadruplexes. Trends in Chemistry, 2020, 2 (2), 123–136. 10.1016/j.trechm.2019.07.002.

(5) Varshney, D.; Spiegel, J.; Zyner, K.; Tannahill, D.; Balasubramanian, S. The Regulation and Functions of DNA and RNA G-Quadruplexes. Nat. Rev. Mol. Cell Biol. 2020, 21 (8), 459–474. 10.1038/s41580-020-0236-x.

(6) Balagurumoorthy, P.; Brahmachari, S. K. Structure and Stability of Human Telomeric Sequence. J. Biol. Chem. 1994, 269 (34), 21858–21869. 10.1016/S0021-9258(17)31882-3.

(7) Robinson, J.; Raguseo, F.; Nuccio, S. P.; Liano, D.; Di Antonio, M. DNA G-Quadruplex Structures: More than Simple Roadblocks to Transcription? Nucleic Acids Res. 2021, 49 (15), 8419–8431. 10.1093/NAR/GKAB609.

(8) Mukherjee, A. K.; Sharma, S.; Chowdhury, S. Non-Duplex G-Quadruplex Structures Emerge as Mediators of Epigenetic Modifications. Trends in Genetics, 2019, 35 (5) 29–144. 10.1016/j.tig.2018.11.001.

(9) Kharel, P.; Becker, G.; Tsvetkov, V.; Ivanov, P. Properties and Biological Impact of RNA G-Quadruplexes: From Order to Turmoil and Back. Nucleic Acids Res. 2020, 48 (22), 12534–12555. 10.1093/NAR/GKAA1126.

(10) Dumas, L.; Herviou, P.; Dassi, E.; Cammas, A.; Millevoi, S. G-Quadruplexes in RNA Biology: Recent Advances and Future Directions. Trends Biochem. Sci. 2021, 46 (4), 270–283. 10.1016/J.TIBS.2020.11.001/ATTACHMENT/356F234B-5586-4E60-B04C-133C43C4CEBD/MMC1.DOCX.

(11) Lyu, K.; Chow, E. Y. C.; Mou, X.; Chan, T. F.; Kwok, C. K. RNA G-Quadruplexes (RG4s): Genomics and Biological Functions. Nucleic Acids Res. 2021, 49 (10), 5426–5450. 10.1093/NAR/GKAB187.

(12) Biffi, G.; Di Antonio, M.; Tannahill, D.; Balasubramanian, S. Visualization and Selective Chemical Targeting of RNA G-Quadruplex Structures in the Cytoplasm of Human Cells. Nat. Chem. 2014, 6 (1), 75. 10.1038/NCHEM.1805.

(13) Biffi, G.; Tannahill, D.; McCafferty, J.; Balasubramanian, S. Quantitative Visualization of DNA G-Quadruplex Structures in Human Cells. Nat. Chem. 2013, 5 (3), 182–186. 10.1038/nchem.1548.

(14) Henderson, A.; Wu, Y.; Huang, Y. C.; Chavez, E. A.; Platt, J.; Johnson, F. B.; Brosh, R. M.; Sen, D.; Lansdorp, P. M. Detection of G-Quadruplex DNA in Mammalian Cells. Nucleic Acids Res. 2014, 42 (2), 860–869. 10.1093/NAR/GKT957.

(15) Chilka, P.; Desai, N.; Datta, B. Small Molecule Fluorescent Probes for G-Quadruplex Visualization as Potential Cancer Theranostic Agents. Molecules. 2019, 24 (4), 752, 10.3390/molecules24040752.

(16) Yuan, J. H.; Shao, W.; Chen, S. Bin; Huang, Z. S.; Tan, J. H. Recent Advances in Fluorescent Probes for G-Quadruplex Nucleic Acids. Biochem. Biophys. Res. Commun. 2020, 531 (1), 18–24. 10.1016/J.BBRC.2020.02.114.

(17) Han, J. N.; Ge, M.; Chen, P.; Kuang, S.; Nie, Z. Advances in G-Quadruplexes-Based Fluorescent Imaging. Biopolymers 2022, 113 (12), e23528. 10.1002/BIP.23528.

(18) Di Antonio, M.; Ponjavic, A.; Radzevičius, A.; Ranasinghe, R. T.; Catalano, M.; Zhang, X.; Shen, J.; Needham, L. M.; Lee, S. F.; Klenerman, D.; Balasubramanian, S. Single-Molecule Visualization of DNA G-Quadruplex Formation in Live Cells. Nat. Chem. 2020, 12, 832–837. 10.1038/s41557-020-0506-4.

(19) Gaigalas, A. K.; Li, L.; Henderson, O.; Vogt, R.; Barr, J.; Marti, G.; Weaver, J.; Schwartz, A. The Development of Fluorescence Intensity Standards. J. Res. Natl. Inst. Stand. Technol. 2001, 106 (2), 381–389. 10.6028/jres.106.015.

(20) Shivalingam, A.; Izquierdo, M. A.; Marois, A. Le; Vyšniauskas, A.; Suhling, K.; Kuimova, M. K.; Vilar, R. The Interactions between a Small Molecule and G-Quadruplexes Are Visualized by Fluorescence Lifetime Imaging Microscopy. Nat. Commun. 2015, 6, 8178. 10.1038/ncomms9178.

(21) Summers, P. A.; Lewis, B. W.; Gonzalez-Garcia, J.; Porreca, R. M.; Lim, A. H. M.; Cadinu, P.; Martin-Pintado, N.; Mann, D. J.; Edel, J. B.; Vannier, J. B.; Kuimova, M. K.; Vilar, R. Visualising G-Quadruplex DNA Dynamics in Live Cells by Fluorescence Lifetime Imaging Microscopy. Nat. Commun. 2021, 12 (1), 1–11. 10.1038/s41467-020-20414-7.

(22) Tseng, T.-Y.; Chien, C.-H.; Chu, J.-F.; Huang, W.-C.; Lin, M.-Y.; Chang, C.-C.; Chang, T.-C. Fluorescent Probe for Visualizing Guanine-Quadruplex DNA by Fluorescence Lifetime Imaging Microscopy. J. Biomed. Opt. 2013, 18 (10), 101309. 10.1117/1.jbo.18.10.101309.

(23) Liu, L.; Liu, W.; Wang, K.; Zhu, B.; Xia, X.; Ji, L.; Mao, Z. Quantitative Detection of G-Quadruplex DNA in Live Cells Based on Photon Counts and Complex Structure Discrimination. Angew. Chemie Int. Ed. 2020, 59 (24), 9719–9726. 10.1002/anie.202002422.

(24) Berezin, M. Y.; Achilefu, S. Fluorescence Lifetime Measurements and Biological Imaging. Chem. Rev. 2010, 110 (5), 2641–2684. 10.1021/cr900343z.

(25) Lu, Y.-J.; Deng, Q.; Hou, J.-Q.; Hu, D.-P.; Wang, Z.-Y.; Zhang, K.; Luyt, L. G.; Wong, W.-L.; Chow, C.-F. Molecular Engineering of Thiazole Orange Dye: Change of Fluorescent Signaling from Universal to Specific upon Binding with Nucleic Acids in Bioassay. ACS Chem. Biol. 2016, 11 (4), 1019–1029. 10.1021/acschembio.5b00987.

(26) Lubitz, I.; Zikich, D.; Kotlyar, A. Specific High-Affinity Binding of Thiazole Orange to Triplex and g-Quadruplex DNA. Biochemistry 2010, 49 (17), 3567–3574. 10.1021/bi1000849.

(27) Long, W.; Lu, Y. J.; Zhang, K.; Huang, X. H.; Hou, J. Q.; Cai, S. Y.; Li, Y.; Du, X.; Luyt, L. G.; Wong, W. L.; Chow, C. F. Boosting the Turn-on Fluorescent Signaling Ability of Thiazole Orange Dyes: The Effectiveness of Structural Modification Site and Its Unusual Interaction Behavior with Nucleic Acids. Dye. Pigment. 2018, 159, 449–456. 10.1016/j.dyepig.2018.07.008.

(28) Zhang, L.; Liu, X.; Lu, S.; Liu, J.; Zhong, S.; Wei, Y.; Bing, T.; Zhang, N.; Shangguan, D. Thiazole Orange Styryl Derivatives as Fluorescent Probes for G-Quadruplex DNA. ACS Appl. Bio Mater. 2020, 3 (5), 2643–2650. 10.1021/acsabm.9b01243.

(29) Jarikote, D. V.; Krebs, N.; Tannert, S.; Röder, B.; Seitz, O. Exploring Base-Pair-Specific Optical Properties of the DNA Stain Thiazole Orange. Chem. – A Eur. J. 2007, 13 (1), 300–310. 10.1002/CHEM.200600699.

(30) Zhao, Z.; Cao, S.; Li, H.; Li, D.; He, Y.; Wang, X.; Chen, J.; Zhang, S.; Xu, J.; Knutson, J. R. Ultrafast Excited-State Dynamics of Thiazole Orange. Chem. Phys. 2022, 553, 111392. 10.1016/J.CHEMPHYS.2021.111392.

(31) Salem, M. A.; Gedik, M.; Brown, A. Two Photon Absorption in Biological Molecules. In Handbook of Computational Chemistry; Springer Netherlands, 2015; pp 1–19. 10.1007/978-94-007-6169-8_47-1.

(32) Helmchen, F.; Denk, W. Deep Tissue Two-Photon Microscopy. Nat. Methods. 2005, 2(12) 932–940. 10.1038/nmeth818.

(33) West, W.; Pearce, S. The Dimeric State of Cyanine Dyes. J. Phys. Chem. 1965, 69 (6), 1894–1903. 10.1021/j100890a019

(34) Ghasemi, J.; Ahmadi, S.; Ahmad, A. I.; Ghobadi, S. Spectroscopic Characterization of Thiazole Orange-3 DNA Interaction. Appl. Biochem. Biotechnol. 2008, 149 (1), 9–22. 10.1007/S12010-007-8124-9/FIGURES/8.

(35) Nygren, J.; Svanvik, N.; Kubista, M. The Interactions between the Fluorescent Dye Thiazole Orange and DNA. Biopolymers 1998, 46 (1), 39–51. 10.1002/(SICI)1097-0282(199807)46:1<39::AID-BIP4>3.0.CO;2-Z.

(36) Biver, T.; Boggioni, A.; Secco, F.; Turriani, E.; Venturini, M.; Yarmoluk, S. Influence of Cyanine Dye Structure on Self-Aggregation and Interaction with Nucleic Acids: A Kinetic Approach to TO and BO Binding. Arch. Biochem. Biophys. 2007, 465 (1), 90–100. 10.1016/J.ABB.2007.04.034.

(37) Das, S.; Purkayastha, P. Modulating Thiazole Orange Aggregation in Giant Lipid Vesicles: Photophysical Study Associated with FLIM and FCS. ACS Omega 2017, 2 (8), 5036–5043. 10.1021/ACSOMEGA.7B00899/ASSET/IMAGES/AO-2017-00899J_M004.GIF.

(38) Klymchenko, A. S.; Emerging Field of Self-Assembled Fluorescent Organic Dye Nanoparticles. J. Nanosci. Lett. 2012, 3, 21. 10.2174/0929867325666180226111716

(39) Masson, T.; Guetta, C. L.; Laigre, E.; Cucchiarini, A.; Duchambon, P.; Teulade-Fichou, M. P.; Verga, D. BrdU Immuno-Tagged G-Quadruplex Ligands: A New Ligand-Guided Immunofluorescence Approach for Tracking G-Quadruplexes in Cells. Nucleic Acids Res. 2021, 49 (22), 12644–12660. 10.1093/NAR/GKAB1166.

(40) Datta, A.; Pollock, K. J.; Kormuth, K. A.; Brosh, R. M. G-Quadruplex Assembly by Ribosomal DNA: Emerging Roles in Disease Pathogenesis and Cancer Biology. Cytogenet. Genome Res. 2021, 161 (6–7), 285–296. 10.1159/000516394.

(41) Laguerre, A.; Hukezalie, K.; Winckler, P.; Katranji, F.; Chanteloup, G.; Pirrotta, M.; Perrier-Cornet, J. M.; Wong, J. M. Y.; Monchaud, D. Visualization of RNA-Quadruplexes in Live Cells. J. Am. Chem. Soc. 2015, 137 (26), 8521–8525. 10.1021/JACS.5B03413/SUPPL_FILE/JA5B03413_SI_001.PDF.

(42) Chen, S. Bin; Hu, M. H.; Liu, G. C.; Wang, J.; Ou, T. M.; Gu, L. Q.; Huang, Z. S.; Tan, J. H. Visualization of NRAS RNA G-Quadruplex Structures in Cells with an Engineered Fluorogenic Hybridization Probe. J. Am. Chem. Soc. 2016, 138 (33), 10382–10385. 10.1021/JACS.6B04799/ASSET/IMAGES/LARGE/JA-2016-04799E_0005.JPEG.

(43) Xu, S.; Li, Q.; Xiang, J.; Yang, Q.; Sun, H.; Guan, A.; Wang, L.; Liu, Y.; Yu, L.; Shi, Y.; Chen, H.; Tang, Y. Thioflavin T as an Efficient Fluorescence Sensor for Selective Recognition of RNA G-Quadruplexes. Sci. Reports 2016 61 2016, 6 (1), 1–9. 10.1038/srep24793.

(44) Zheng, B.-X.; Long, W.; She, M.-T.; Wang, Y.; Zhao, D.; Yu, J.; Siu-Lun Leung, A.; Hin Chan, K.; Hou, J.; Lu, Y.-J.; Wong, W.-L.; Zheng, B.; Long, W.; Yu, J.; Leung, A.; Chan, K. H.; Wong, W.; Wang, Y.; She, M.; Lu, Y.; Zhao, D.; Hou, J. A Cytoplasm-Specific Fluorescent Ligand for Selective Imaging of RNA G-Quadruplexes in Live Cancer Cells. Chem. – A Eur. J. 2023, 29 (34) e202300705. 10.1002/CHEM.202300705.

(45) Ivanov, P.; O’Day, E.; Emara, M. M.; Wagner, G.; Lieberman, J.; Anderson, P. G-Quadruplex Structures Contribute to the Neuroprotective Effects of Angiogenin-Induced TRNA Fragments. Proc. Natl. Acad. Sci. U. S. A. 2014, 111 (51), 18201–18206. 10.1073/PNAS.1407361111/-/DCSUPPLEMENTAL.

(46) Bhattacharyya, D.; Arachchilage, G. M.; Basu, S. Metal Cations in G-Quadruplex Folding and Stability. Front. Chem. 2016, 4, 38. 10.3389/FCHEM.2016.00038.

(47) del Villar-Guerra, R.; Trent, J. O.; Chaires, J. B. G-Quadruplex Secondary Structure from Circular Dichroism Spectroscopy. Angew. Chem. Int. Ed. Engl. 2018, 57 (24), 7171. 10.1002/ANIE.201709184.

(48) Horobin, R. W.; Stockert, J. C.; Rashid-Doubell, F. Uptake and Localisation of Small-Molecule Fluorescent Probes in Living Cells: A Critical Appraisal of QSAR Models and a Case Study Concerning Probes for DNA and RNA. Histochem. Cell Biol. 2013, 139 (5), 623–637. 10.1007/S00418-013-1090-0/TABLES/6.

(49) Delic, J.; Coppey, J.; Magdelenat, H.; Coppey-Moisan, M. Impossibility of Acridine Orange Intercalation in Nuclear DNA of the Living Cell. Exp. Cell Res. 1991, 194 (1), 147–153. 10.1016/0014-4827(91)90144-J.

(50) Tramier, M.; Kemnitz, K.; Durieux, C.; Coppey, J.; Denjean, P.; Pansu, R. B.; Coppey-Moisan, M. Restrained Torsional Dynamics of Nuclear DNA in Living Proliferative Mammalian Cells. Biophys. J. 2000, 78 (5), 2614–2627. 10.1016/S0006-3495(00)76806-8.

(51) Monchaud, D.; Allain, C.; Bertrand, H.; Smargiasso, N.; Rosu, F.; Gabelica, V.; De Cian, A.; Mergny, J. L.; Teulade-Fichou, M. P. Ligands Playing Musical Chairs with G-Quadruplex DNA: A Rapid and Simple Displacement Assay for Identifying Selective G-Quadruplex Binders. Biochimie. 2008, 90 (8), 1207–1223. 10.1016/J.BIOCHI.2008.02.019.

(52) De Cian, A.; DeLemos, E.; Mergny, J. L.; Teulade-Fichou, M. P.; Monchaud, D. Highly Efficient G-Quadruplex Recognition by Bisquinolinium Compounds. J. Am. Chem. Soc. 2007, 129 (7), 1856–1857. 10.1021/JA067352B/SUPPL_FILE/JA067352BSI20070110_014417.PDF.

(53) Abd Karim, N. H.; Mendoza, O.; Shivalingam, A.; Thompson, A. J.; Ghosh, S.; Kuimova, M. K.; Vilar, R. Salphen Metal Complexes as Tunable G-Quadruplex Binders and Optical Probes. RSC Adv. 2013, 4 (7), 3355–3363. 10.1039/C3RA44793F.

(54) Reed, J. E.; Arnal, A. A.; Neidle, S.; Vilar, R. Stabilization of G-Quadruplex DNA and Inhibition of Telomerase Activity by Square-Planar Nickel(II) Complexes. J. Am. Chem. Soc. 2006, 128 (18), 5992–5993. 10.1021/JA058509N/SUPPL_FILE/JA058509NSI20060320_012503.PDF.

(55) Kosiol, N.; Juranek, S.; Brossart, P.; Heine, A.; Paeschke, K. G-Quadruplexes: A Promising Target for Cancer Therapy. Mol. Cancer. 2021, 20 (1), 40. 10.1186/s12943-021-01328-4.

(56) Wang, E.; Thombre, R.; Shah, Y.; Latanich, R.; Wang, J. G-Quadruplexes as Pathogenic Drivers in Neurodegenerative Disorders. Nucleic Acids Res. 2021, 49 (9), 4816–4830. 10.1093/NAR/GKAB164.

(57) Zheng, K. W.; Zhang, J. Y.; He, Y. De; Gong, J. Y.; Wen, C. J.; Chen, J. N.; Hao, Y. H.; Zhao, Y.; Tan, Z. Detection of Genomic G-Quadruplexes in Living Cells Using a Small Artificial Protein. Nucleic Acids Res. 2020, 48 (20), 11706–11720. 10.1093/nar/gkaa841.

(58) Vummidi, B. R.; Alzeer, J.; Luedtke, N. W. Fluorescent Probes for G-Quadruplex Structures. ChemBioChem 2013, 14 (5), 540–558. 10.1002/CBIC.201200612.

(59) Guo, J. U.; Bartel, D. P. RNA G-Quadruplexes Are Globally Unfolded in Eukaryotic Cells and Depleted in Bacteria. Science 2016, 353 (6306). 10.1126/SCIENCE.AAF5371.

(60) Raguseo, F.; Tanase, D.; Malouf, L.; Sanchez, R. M. R.; Elani, Y.; Michele, L. Di; Antonio, M. Di. The C9Orf72 Hexanucleotide Repeat Expansion Transcript Forms Insoluble Aggregates Facilitated by Multimolecular G-Quadruplex Structures. bioRxiv 2023, 10.1101/2023.01.31.526399.

(61) Turner, M.; Danino, Y. M.; Barshai, M.; Yacovzada, N. S.; Cohen, Y.; Olender, T.; Rotkopf, R.; Monchaud, D.; Hornstein, E.; Orenstein, Y. RG4detector, a Novel RNA G-Quadruplex Predictor, Uncovers Their Impact on Stress Granule Formation. Nucleic Acids Res. 2022, 50 (20), 11426. 10.1093/NAR/GKAC950.

(62) Zhang, Y.; Yang, M.; Duncan, S.; Yang, X.; Abdelhamid, M. A. S.; Huang, L.; Zhang, H.; Benfey, P. N.; Waller, Z. A. E.; Ding, Y. G-Quadruplex Structures Trigger RNA Phase Separation. Nucleic Acids Res. 2019, 47 (22), 11746–11754. 10.1093/nar/gkz978.

