## Supplementary information for "Cellular visualization of G-quadruplex RNA via fluorescence lifetime imaging microscopy"

### 1. Synthetic chemistry

Compound **1**, **2** and **3** were prepared following previously reported synthetic procedures.^1^ **TOR-G4** was synthesized from **3** as shown in Scheme 1.

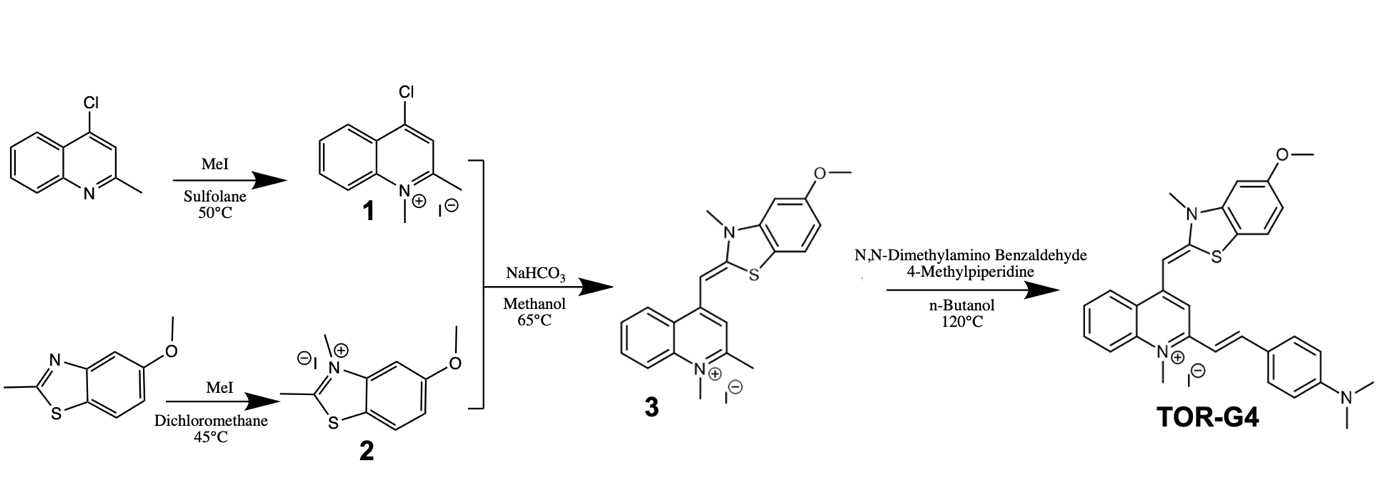

**Scheme 1** - Reaction scheme for the synthesis of **TOR-G4**.

**Synthesis of compound 1**: 4-chloro-2-methylquinoline (0.6 mL, 2.8 mmol) and iodomethane (1.1 mL, 16.7 mmol) were refluxed in sulfolane (10 mL) at 50 ºC overnight. A colour change of pale yellow to dark purple was observed. The solution was cooled to room temperature and anhydrous diethyl ether (7.5 mL) was added and stirred for 30 min. A solid dark purple precipitate formed, which was filtered and dried under vacuum. To remove remaining sulfolane, the product was re-dissolved in chloroform (~5 mL), precipitated with the addition of anhydrous diethyl ether (~5 mL) and dried under vacuum (516 mg, 58% yield, **1**). ^1^H NMR spectrum (Figure S1) was recorded in DMSO-d_6_ at 400MHz (δ in ppm): 8.67 (d, J=9.0 Hz, 1H_d_), 8.56 (dd, J= 8.4, 1.3 Hz, 1H_a_), 8.54 (s, 1H_g_), 8.34 (m, 1H_b_), 8.13 (t, J=9 Hz, 1H_c_), 4.43 (s, 3H_e_), 3.06 (s, 3H_f_).

**Synthesis of compound 2:** 5-methoxy-2-methylbenzo[*d*]thiazole (500 mg, 2.77 mmol) was dissolved in dichloromethane (10 mL) and refluxed with iodomethane (3.5 mL, 55.8 mmol) at 45 ºC overnight. The solution was cooled to room temperature and anhydrous diethyl ether (10 mL) was added and stirred for 30 min. A fine white precipitate was formed which was pelleted by centrifugation and dried under vacuum to yield the product (382 mg, 43%, **2**). ^1^H NMR spectrum (Figure S2) was recorded in DMSO-d_6_ at 400 MHz (δ in ppm): 8.30 (d, J = 9.0 Hz, 1H_a_), 7.79 (d, J = 2.3 Hz, 1H_d_), 7.44 (dd, J = 9.0, 2.3 Hz, 1H_b_), 4.18 (s, 3H_e_), 3.98 (s, 3H_c_), 3.14 (s, 3H_f_).

**Synthesis of compound 3**: **1** (80 mg, 0.25 mmol) and **2** (80 mg, 0.25 mmol) were dissolved in methanol (10 mL), sodium bicarbonate solution (0.5 M, 1 mL) and saturated potassium iodide solution (2 mL) and refluxed at 65 ºC for 2h. The solution was cooled to room temperature and stirred in water for 20 min before being filtered and dried under vacuum. The product was a bright orange-red solid (22 mg, 19%, **3**). ^13^C NMR (500 MHz, DMSO, Figure S3), δ (ppm): 160.2 (C4), 160.1 (C6), 153.9 (C13), 147.5 (C9), 141.9 (C15), 139.0 (C20), 133.0 (C1), 126.3 (C2), 125.3 (C3), 123.4 (C19), 123.3 (C17), 118.2 (C11), 114.9 (C6), 112.0 (C18), 110.4 (C8), 98.0 (C16), 87.0 (C12), 56.0 (C5), 37.0 (C21), 33.8 (C14), 22.8 (C7). ^1^H NMR (400 MHz, DMSO-d_6_ Figure S4), δ (ppm): 8.77 (d, J = 8.3 Hz, 1H_d_), 8.18 (d, J = 8.7 Hz, 1H_a_), 8.04 – 7.95 (m, 1H_b_), 7.89 (d, J = 8.7 Hz, 1H_m_), 7.75 (t, J = 7.5 Hz, 1H_c_), 7.35 (d, J = 2.3 Hz, 1H_j_), 7.29 (s, 1H_g_), 7.02 (dd, J = 8.7, 2.3 Hz, 1H_l_), 6.86 (s, 1H_h_), 4.07 (s, 3H_e_), 4.00 (s, 3H_k_), 3.91 (s, 3H_i_), 2.87 (s, 3H_f_).

**Synthesis of (TOR-G4)**:

**3** (18 mg, 0.038 mmol) was dissolved in n-butanol (7.5 mL) and 4-methyl piperidine (0.5 mL) and stirred at room temperature for 15 min. 4-dimethylamino benzaldehyde (11 mg, 0.076 mmol) was added and the solution was refluxed to 120 ºC for 3h. A dark brown precipitate formed which was filtered, washed with n-butanol and dried in vacuo to yield the final product (5 mg, 22%, **TOR-G4**). ^13^C NMR (500 MHz, DMSO, Figure S5), δ (ppm): 160.0 (C4), 159.4 (C6), 152.7 (C13), 151.8 (C26), 146.6 (C9), 142.1 (C25), 142.0 (C15), 139.1 (C20), 132.9 (C1), 130.4 (C24), 126.1 (C2), 125.0 (C3), 123.5 (C19), 123.4 (C17), 122.7 (C22), 118.4 (C11), 114.8 (C6), 114.7 (C23), 111.7 (C7), 111.5 (C18), 107.5 (C8), 97.8 (C16), 87.3 (C12), 55.9 (C5), 37.7 (C21), 33.6 (C14). (NMe_2_ in solvent peak). ^1^H NMR (400 MHz, DMSO-d_6_, Figure S6), δ (ppm): 8.68 (d, J = 9.0 Hz, 1H_d_), 8.09 (d, J = 9.5 Hz, 1H_a_), 7.96 – 7.86 (m, 2H_b/l_), 7.77 (d, J = 8.9 Hz, 2H_o_), 7.69 (dd, J = 8.2, 6.8 Hz, 1H_c_), 7.60 (d, J = 15.6 Hz, 1H_n_), 7.54 (s, 1H_f_), 7.41 (d, J = 15.6 Hz, 1H_m_), 7.24 (d, J = 2.3 Hz, 1H_i_), 6.98 (dd, J = 8.7, 2.3 Hz, 1H_k_), 6.79 (m, 3H_g/p_), 4.10 (s, 3H_e_), 3.94 (s, 3H_j_), 3.89 (s, 3H_h_), 3.05 (s, 6H_q_). LC-MS (Figure S7) found a 480 Da ES+ species, which corresponds to the expected mass of **TOR-G4**.

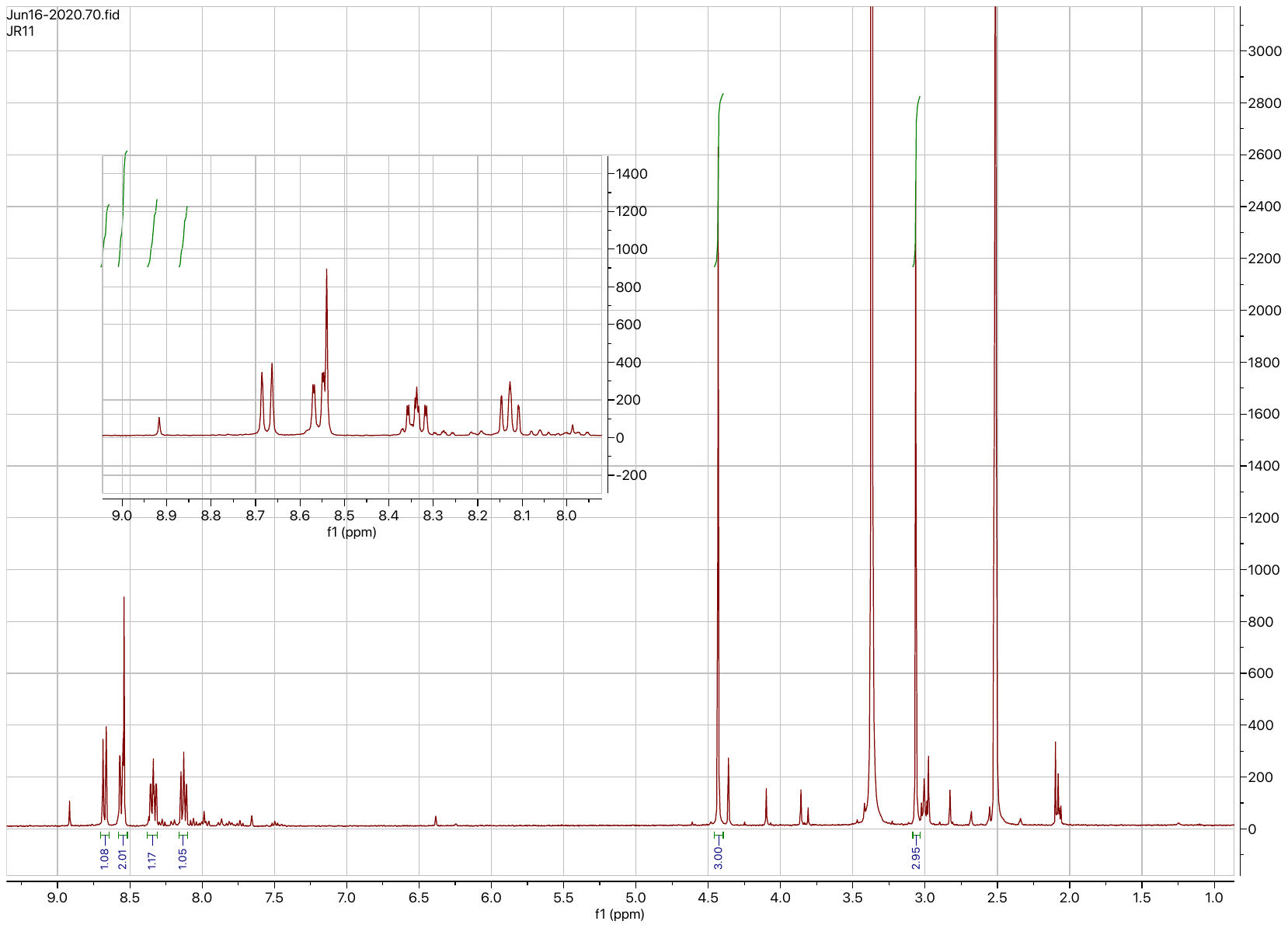

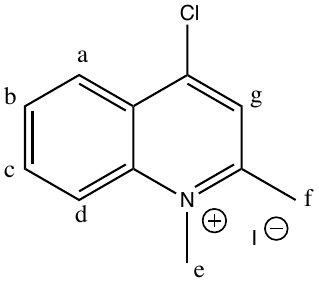

**Figure S1** – ^1^H NMR spectrum of **1** in DMSO-d^6^.

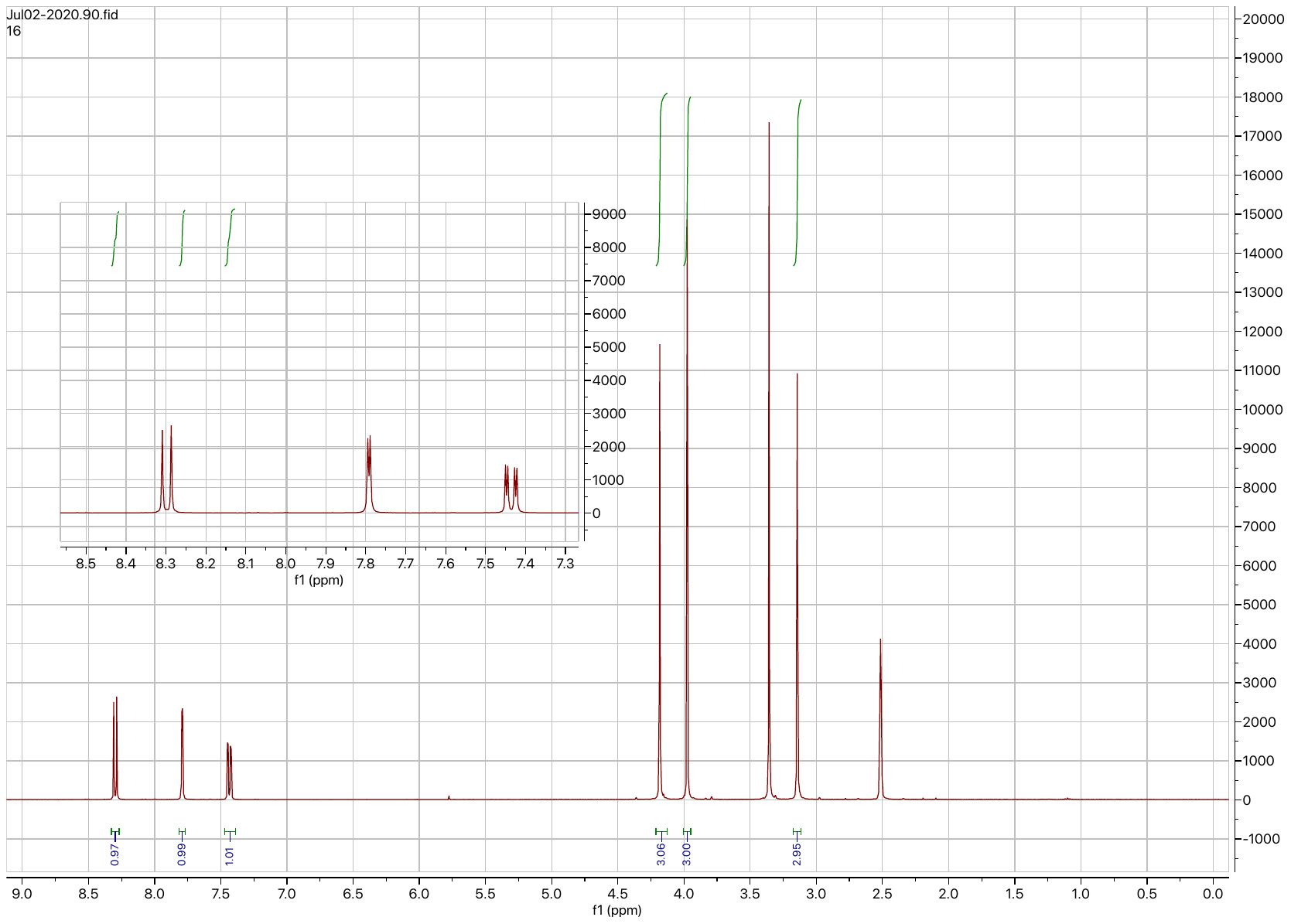

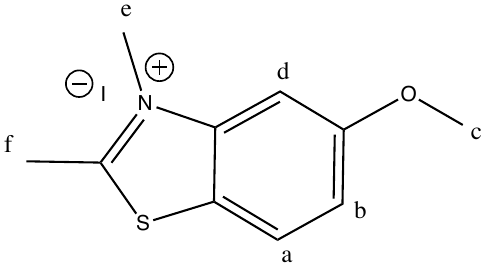

**Figure S2** – ^1^H NMR spectrum of **2** in DMSO-d^6^.

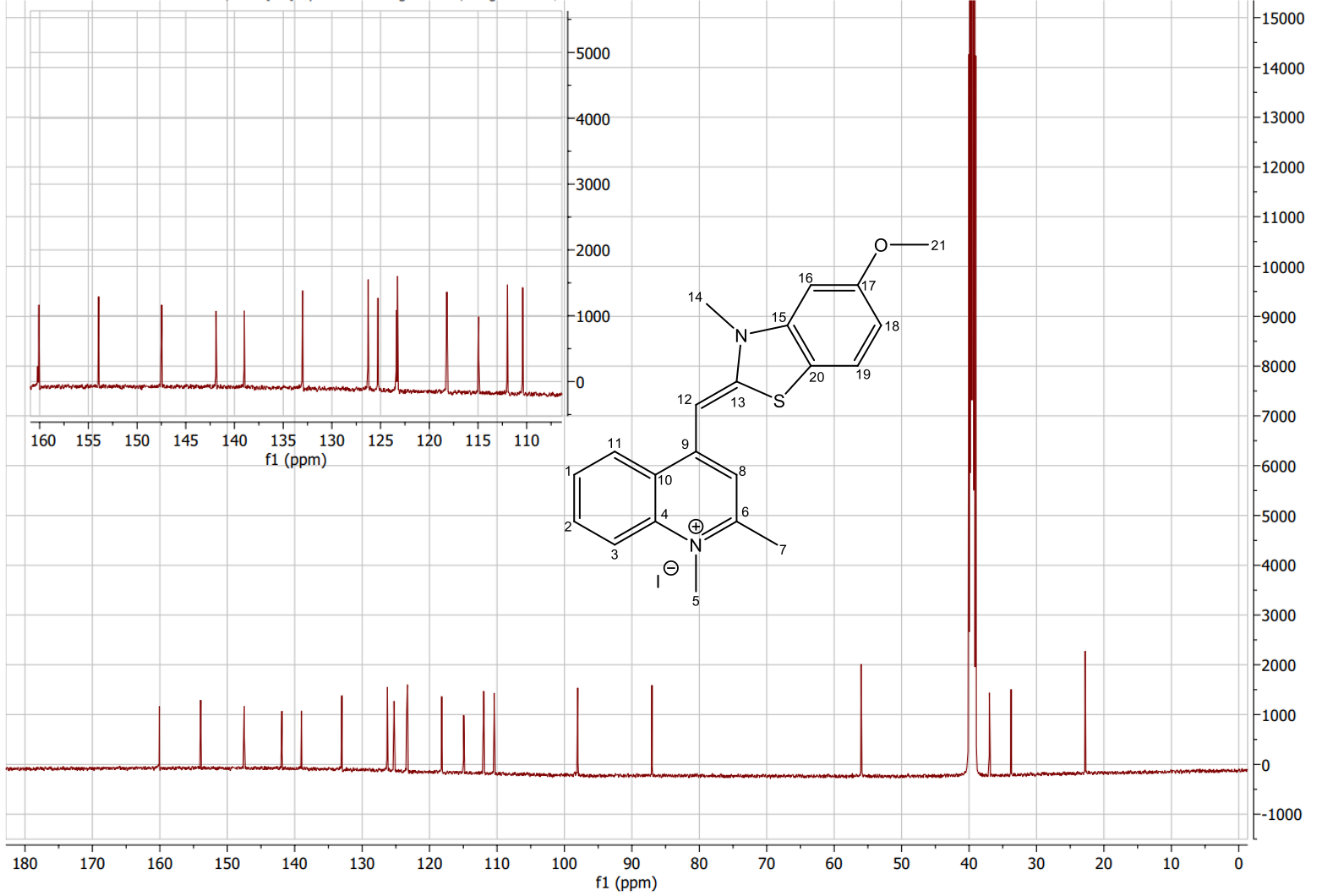

**Figure S3** – ^13^C NMR spectrum of **3** in DMSO-d^6^.

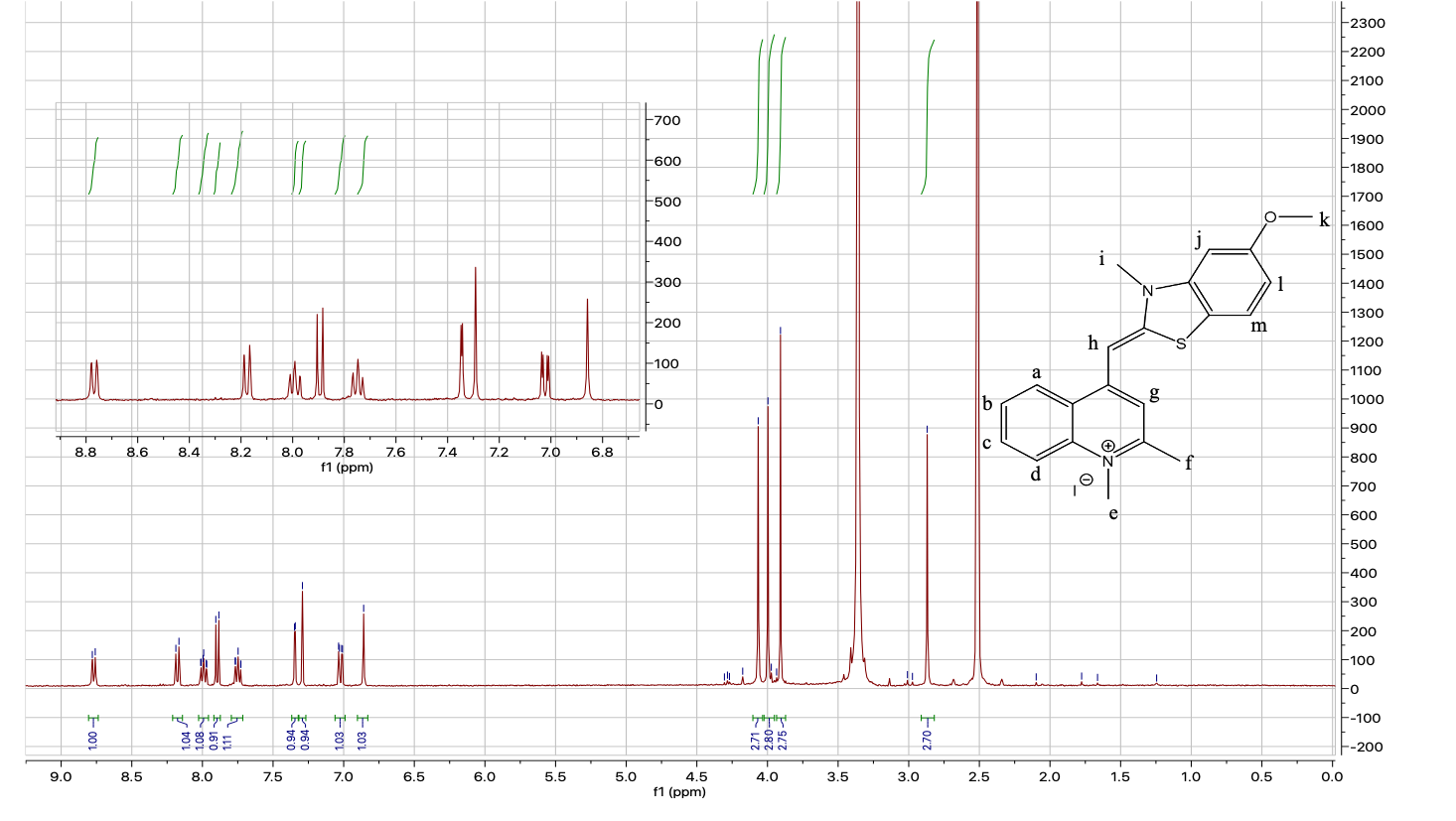
**Figure S4** – ^1^H NMR spectrum of **3** in DMSO-d^6^

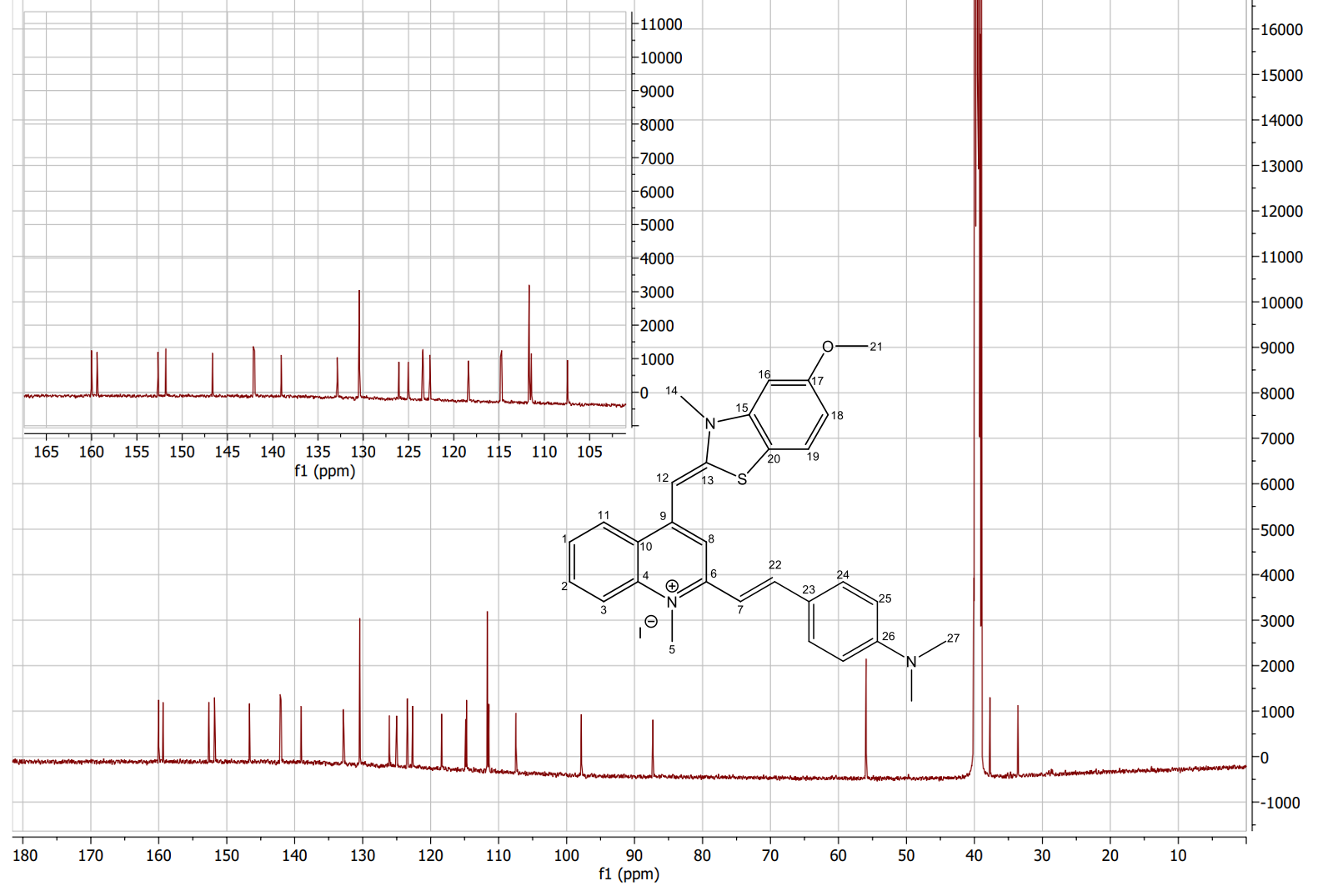

**Figure S5** – ^13^C NMR spectrum of **TOR-G4** in DMSO-d^6^

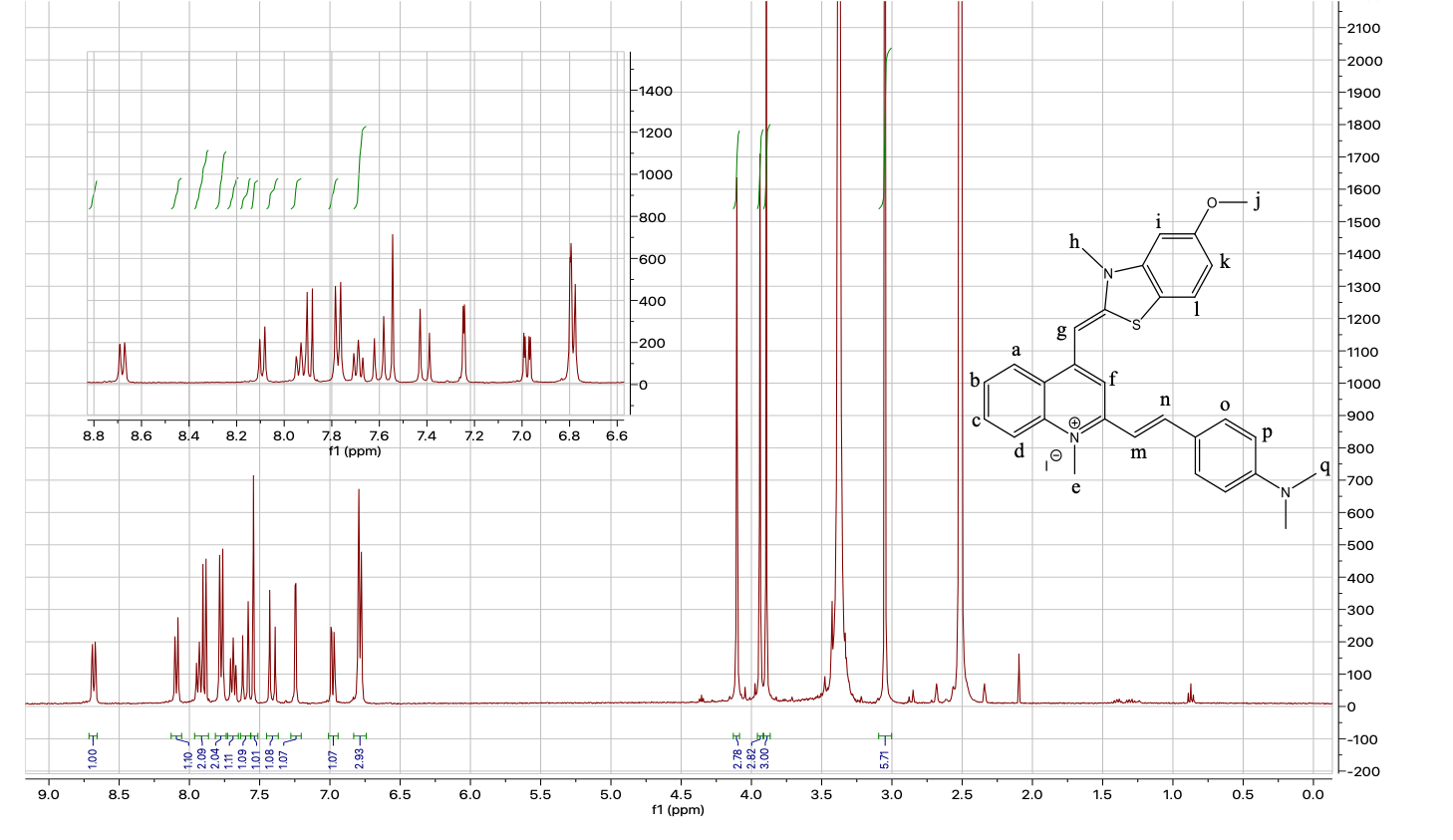

**Figure S6** – ^1^H NMR spectrum of **TOR-G4** in DMSO-d^6^.

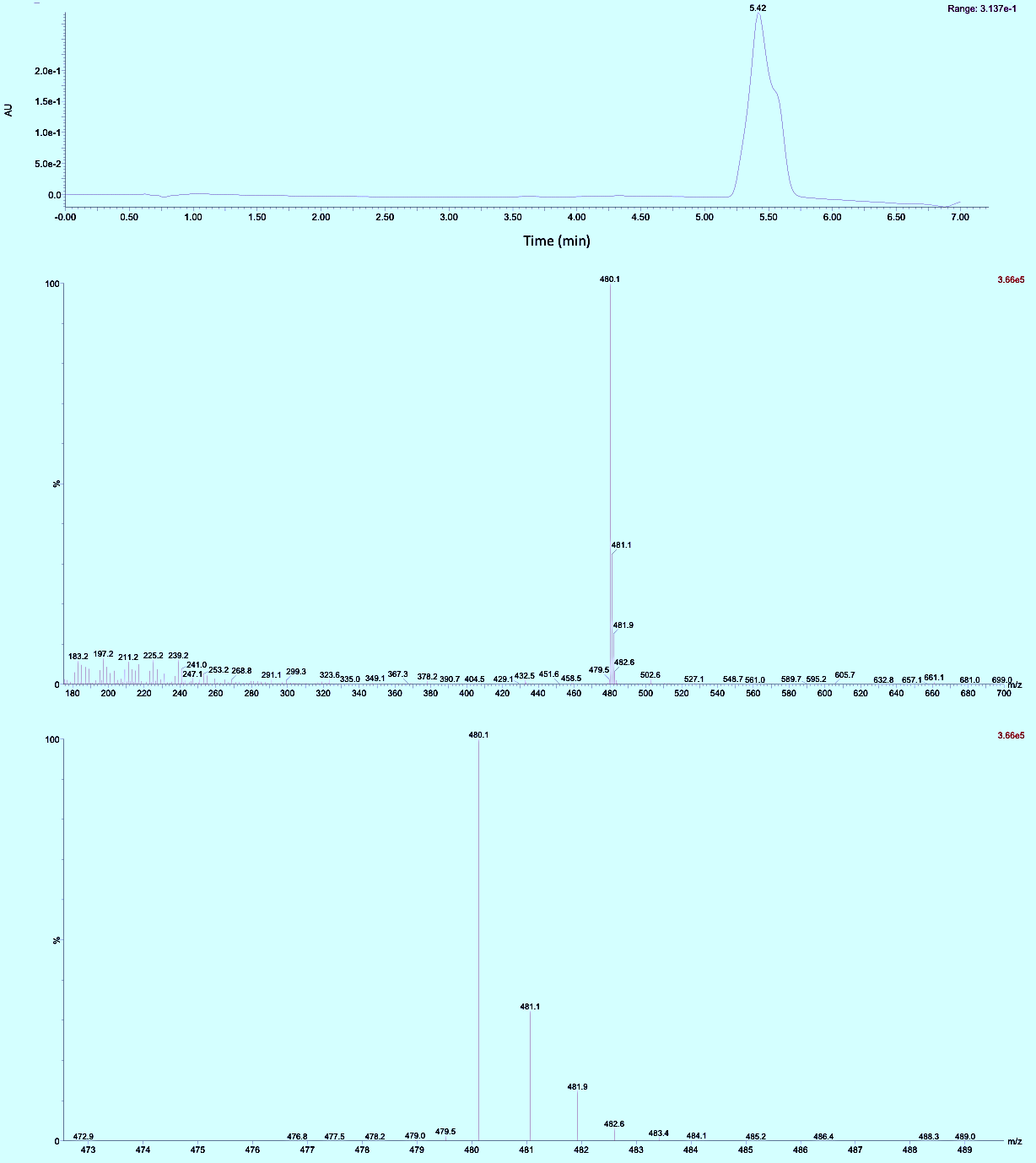

**Figure S7** – LC-MS of **TOR-G4**.

### 2. Photophysical characterization

**Table S1** – Photophysical properties of TOR-G4 in aqueous buffer and in the presence of a 10-fold excess of G4 (*c-MYC*).

| Condition | Excitation max 1 (nm) | Excitation max 2 (nm) | Emission max 1 (nm)^a^ | Emission max 2 (nm)^a^ | Extinction coefficient^b^ (M^-1^ cm^-1^) | Quantum yield  $\boldsymbol{\phi}$_f_ |
| --- | --- | --- | --- | --- | --- | --- |
| TOR-G4 only | 453 | 506 | 710 | - | 20,356 | 0.02^c^ |
| TOR-G4 + 10x G4 (c-MYC) | 488 | 570 | 540 | 660 | 20,026 | 0.1^d^ |

^a^ emission measured after excitation at 470 nm

^b^ extinction coefficient measured at 470 nm

^c^ quantum yield measured at 710 nm. Cresyl violet ($\phi$_f_ = 0.54 in methanol) was used as a standard for quantum yield experiments.

^d^ quantum yield measured for 660 nm species

DNA and RNA oligos were purchased from IDT, with standard desalting purification and used as received. All reagents and solvents (spectroscopic grade) were purchased from Sigma.

All experiments were performed in Tris-HCl buffer (10 mM, pH 7.4) supplemented with 100 mM KCl (or 100 mM LiCl for G4 lithium experiments).

**Table S2**– List of all nucleic acid sequences characterized

| Name | Structure | Sequence |
| --- | --- | --- |
| TRF2 | G4 RNA – parallel | CGGGAGGGCGGGGAGGGC |
| BCL2 | G4 RNA – parallel | AGGGGGCCGUGGGGUGGGAGCUGGGG |
| NRAS | G4 RNA – parallel | GGGAGGGGCGGGUCUGGG |
| c-MYC | G4 DNA – parallel | TGAGGGTGGGTAGGGTGGGTAA |
| HTG4 | G4 DNA – hybrid | TTGGGTTAGGGTTAGGGTTAGGGA |
| HRAS | G4 DNA – antiparallel | TCGGGTTGCGGGCGCAGGGCACGG |
| ssRNA-1 | Single stranded RNA | UGAGCUUAAUUGUAUAUAUUCG |
| ssRNA-2 | Single stranded RNA | CAAUUGUAUAUAUUCG |
| hairpin | Hairpin RNA | CAGUACAGAUCUGUACUG |
| Cov-bulge | Hairpin with symmetric bulge | AGUGCUUCAGUCAGCUGAUGCACU |
| Ucu-bulge | Hairpin with asymmetric bulge | GGCCAGAUCUGAGCCUGGGAGCUCUCUGGCC |
| yeast totRNA | All RNA extracted from yeast (sigma) | NA |
| tRNA | tRNA extract from yeast (sigma) | NA |
| Duplex DNA | Duplex DNA extracted from calf thymus (sigma) | NA |

Absorption spectra were recorded on an Agilent 8453 UV-Visible spectrophotometer across a range of 190-1100 nm in 1 nm intervals. Emission spectra were recorded on a Fluoromax4-spectrofluorimeter (Horiba Jobin-Yvon) with excitation at 470 nm. Excitation spectra were recorded on a Fluoromax4-spectrofluorimeter (Horiba Jobin-Yvon) with detection at 660 nm and excitation from 345 – 650 nm or detection at 540 nm and excitation from 275 – 530 nm.

*In vitro* fluorescence lifetime measurements were made via time correlated single photon counting using a DeltaFlex modular lifetime system (Jobin-Ybon, Horiba), coupled to a 467 nm NanoLED diode laser (pulse width <200 ps, Horiba) as an excitation source. To measure the instrument response function (IRF), prompt measurements were made using a dilute LUDOX solution at the laser excitation wavelength. Emission was monitored at 540 nm with a 32 nm bandpass, across a 100 ns time scale **(**split between 4096 time bins) until 10**,**000 counts were reached in the maximum. A long-pass filter at 495 nm was used to block the detection of scattered excitation light during the measurements. The time resolved decays were fitted to a bi-exponential decay function (equation 1) with deconvolution from the IRF, using Horiba DAS6 lifetime analysis software. Intensity-weighted average lifetimes ($\tau$_w_) were calculated from the individual amplitudes (A) and lifetimes ($\boldsymbol{\tau)}$ values extracted from each decay according to equation 2.

(1) Y(t) = A_1_e^-t/τ1^ + A_2_e^-t/τ2^
(2) $\tau_{W}=\frac{{{\sum A_{i}\tau}_{i}}^{2}}{\sum{A_{i}\tau}_{i}}$

Fluorescence switch-on was calculated by dividing the emission of **TOR-G4** (at 540 nm) in the presence of a given nucleic acid by that of the probe alone in aqueous buffer.

DNA/RNA binding titrations were performed by recording the fluorescence intensity of **TOR-G4** (2 $\mu$M) at 540 nm following 470 nm excitation, with varying concentrations of each nucleic acid (0-150 $\mu$g/mL). Fluorescence intensity was normalized for absorbance of each solution at 470 nm and plotted against nucleic acid concentration. The resulting curves were fitted using Graphpad prism one site –total model.

The G4/totRNA titration was performed by measuring the lifetime of **TOR-G4** (2 $\mu$M) bound to yeast totRNA (100 $\mu$g/mL) and increasing amounts of G4 RNA (TRF-2). The subsequent response curve was then fit in Graphpad prism using the [agonist] vs response – variable slop (4 parameters) model.

G4 displacement assays were performed by addition of increasing concentrations (0.25 - 2 $\mu$M) of Ni-Salphen (synthesized according to literature procedure)^2^ or PhenDC3 (Merck, >97% purity) to **TOR-G4** (2 $\mu$M) bound to NRAS G4, hairpin or yeast totRNA sequences (100 $\mu$g/mL). Fluorescence intensity at 540 nm were recorded following 470 nm excitation; time resolved fluorescence decays were recorded as described above.

### 3. Molecular modelling

Molecule geometry was optimized with Gaussian 09 software using DFT calculations utilizing the B3LYP functional and 6-31g basis set. The duplex DNA structure was taken from the RCSB protein data bank (pdb108D) based on the solution NMR structure of a DNA sequence in complex with homo-dimeric thiazole orange (TOTO).^3^ The G-quadruplex structure was taken from the protein data bank structure of the *c-Myc* promoter in complex with a quindoline ligand (pdb2L7V).^4^ The dimensions of the active site box were selected using AutoDock tools and were set to cover the entirety of the DNA structure. AutoDock Vina was used to run molecular dynamic measurements using default settings, energy range=9 and exhaustiveness=9.^5^ Output files were visualized with PyMOL.

### 4. Cellular characterization

U2OS cells (ECACC) were plated (2x10^4^ cells per well, 250 $\mu$L, 0.8 cm^2^) 24 hours before each experiment and grown in DMEM media (Gibco) supplemented with 10% FBS. For fixed cell experiments, cells were washed three times with ice cold PBS before fixation with 4% paraformaldehyde for 10 min. Cells were then washed again three times with ice cold PBS, before probe staining.

Cell lysate was extracted from 1x10^7^ U2OS cells by first scraping and pelleting cells. Cell pellets were then lysed with ice-cold Chromatrap hypotonic buffer (500 $\mu$L, 10 min). Further nuclear lysis occurred via treatment on ice with Chromatrap lysis buffer (100 $\mu$L, 10 min). Lysate was then diluted in nuclease-free water (1 mL) to yield a final concentration of 90 ug/mL of RNA and 30 ug/mL of DNA quantified using a qubit 4 fluorimeter and the qubit RNA and DNA broad range kits. Nucleic acids were removed by treating lysate with benzonase (1,000 units, 24 hours, 37 °C, Millipore).

RNA was extracted from U2OS cells (4 x10^6^) using Qiagen’s RNeasy extraction kit, following the manufacturer’s instructions. Briefly: adherent cells were trypsinized, counted and aliquoted. Cell pellets were then lysed with the RNeasy lysis buffer and ran through Qiagen’s QIAshedder homogenizing columns. Cell lysate was then washed with RNeasy wash buffer and an on-column DNase I digestion was performed, followed by RNA purification according to the kit’s instructions and RNA elution with RNase-free water (50 $\mu$L). The final concentration of RNA obtained was measured on a NanoDrop One UV-Vis spectrophotometer.

To measure cellular toxicity of the probe, MTS assays were conducted using the Promega CellTiter 96 Aqueous One Solution Cell Proliferation Assay kit, following manufactures instructions. Cells were incubated with **TOR-G4** over 6 hours at varying concentrations from 5-2,000 nM. Absorbance of the MTS reagent was then measured at 490 nm. The experiment was conducted in triplicate and survival curves were fit using the GraphPad Prism Inhibitor vs. response - Variable slope model.

Confocal images were acquired using a Leica SP5 II confocal microscope after incubation of fixed U2OS cells with **TOR-G4** (5 $\mu$M, no washing) for 2 hours. Images were taken with a 100× oil immersion objective (correction collar, NA = 1.2, Leica) after excitation using an internal Ar+ laser at 514 nm and detection at 550-700 nm.. For co-staining experiments, DNA was stained with DAPI (10 $\mu$M, 30 min, two-photon excitation at 760 nm, detection = 400-500 nm) and RNA with SYTO RNASelect (500 nM, 30 min, excitation = 514 nm, detection = 550-700 nm).

FLIM images were obtained on a Leica SP5 II confocal microscope coupled to a TCSPC module (Becker & Hickl GmbH) following excitation with a pulsed diode laser at 477 nm (Becker & Hickl GmbH, 20 MHz). For two-photon excitation FLIM images, a femtosecond Ti:sapphire laser (Coherent, 80 MHz) at 760 nm was used. Fluorescence was collected in a 550-700 nm window with a PMC-100-1 photomultiplier tube detector (Hamamtsu) for 500 s. Images were acquired at 256 x 256-pixel resolution. High resolution zoomed images were acquired with 516 x 516 pixels, this resulted in larger distinctions in lifetime being seen between nucleoli and rest of the nucleus and a corresponding change in the pseudo pixel coloring.

FLIM images were analyzed with FLIMfit software (Imperial College London)^6^: whole cells were manually segmented and fluorescence decays were fitted pixel-wise to a bi-exponential decay function using the maximum likelihood algorithm, with deconvolution from the IRF. Scattering of light and peak offset were fitted locally using FLIMfit default settings. The reported cellular lifetimes are the intensity-weighted average fluorescence lifetimes as calculated in equation 2.

Nuclease experiments were performed by monitoring **TOR-G4** fluorescence following the incubation of fixed cells with either DNase I (200 units/well, Qiagen), RNase H (0.2 U/$\mu$L, New England BioLabs), Ambion RNase A (0.1 µg/$\mu$L, Invitrogen) or RNase T1 (2 U/$\mu$L, Life Technologies) at 37 °C for 30 mins prior to probe incubation.

Transcriptional inhibition was achieved by incubation of cells with DRB (5,6-dichloro-1-β-d-ribofuranosyl-1h-benzimidazole, 100 $\mu$M, Cambridge bioscience) for 1 hour prior to imaging.

For the G4 displacement assay, Ni-Salphen or PhenDC3 (1 µM) were added to cells following probe incubation and imaged by FLIM over approximately 7 hours.

RNA transfection was performed with lipofectamine 2000 (Invitrogen) following the manufacturer’s instructions: G4 RNA (NRAS, 1 µg) or hairpin RNA (1 µg) was incubated with lipofectamine at a 3:1 ratio of lipofectamine:RNA in Optimem media (Gibco) for 15 min. The lipofectamine/RNA mixture was then incubated with cells for 6 hours, before cell fixation and imaging with **TOR-G4**.

Statistical significance of perturbation experiments was assessed with the Welch t-test in Graphpad prism.

### 5. Supplementary figures

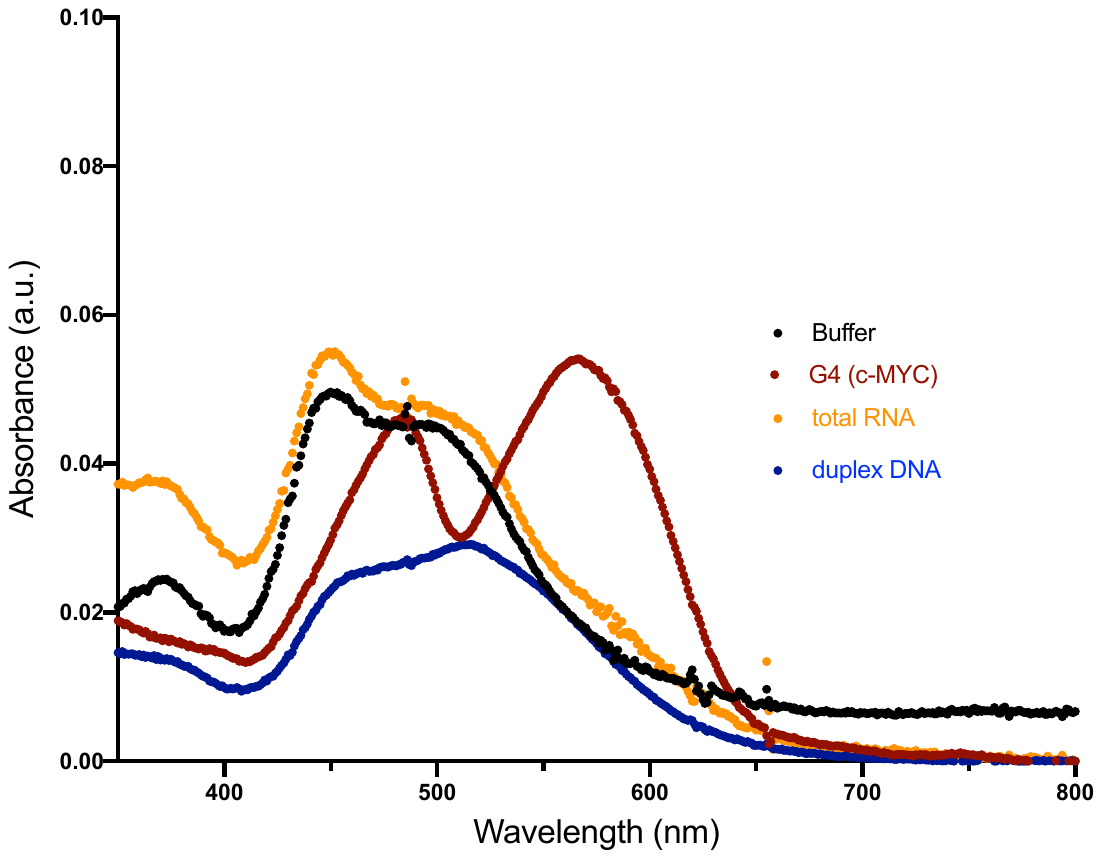

**Figure S8 –** Absorbance spectra of **TOR-G4** (2 $\mu$M) in aqueous buffered solutions and bound to G4, total RNA and duplex DNA sequences (300 $\mu$g/mL). A clear spectral change is visible upon interaction of **TOR-G4** with G4 DNA.

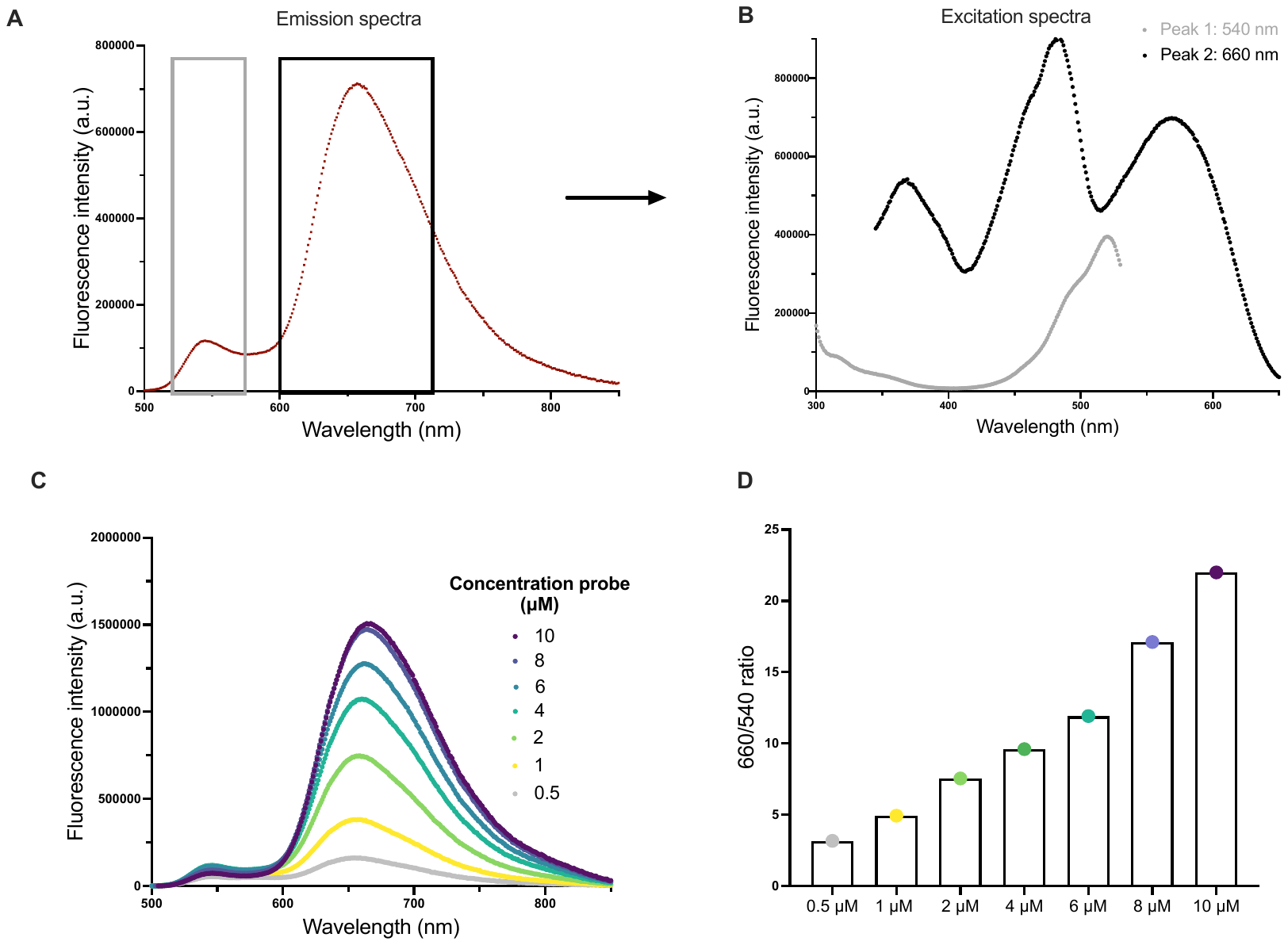
**Figure S9** – Understanding **TOR-G4** aggregation - A) Emission and B) excitation spectra of **TOR-G4** (2 $\mu$M) bound to G4 DNA (c-MYC, 100 $\mu$g/mL); A clear change in the excitation spectra is seen for the two peaks (cf Fig S8, absorption spectra of **TOR-G4)** C) Emission spectra recorded for increasing concentrations of **TOR-G4** (0.5-10 $\mu$M) in the presence of c-MYC DNA (100 $\mu$g/mL) and D) corresponding ratios of 660/540 nm band intensity. The significant increase in the intensity of the 660 nm band with increased **TOR-G4** concentration indicate that this band corresponds to **TOR-G4** aggregation, whilst the 540 nm band is likely due to **TOR-G4** monomer emission.

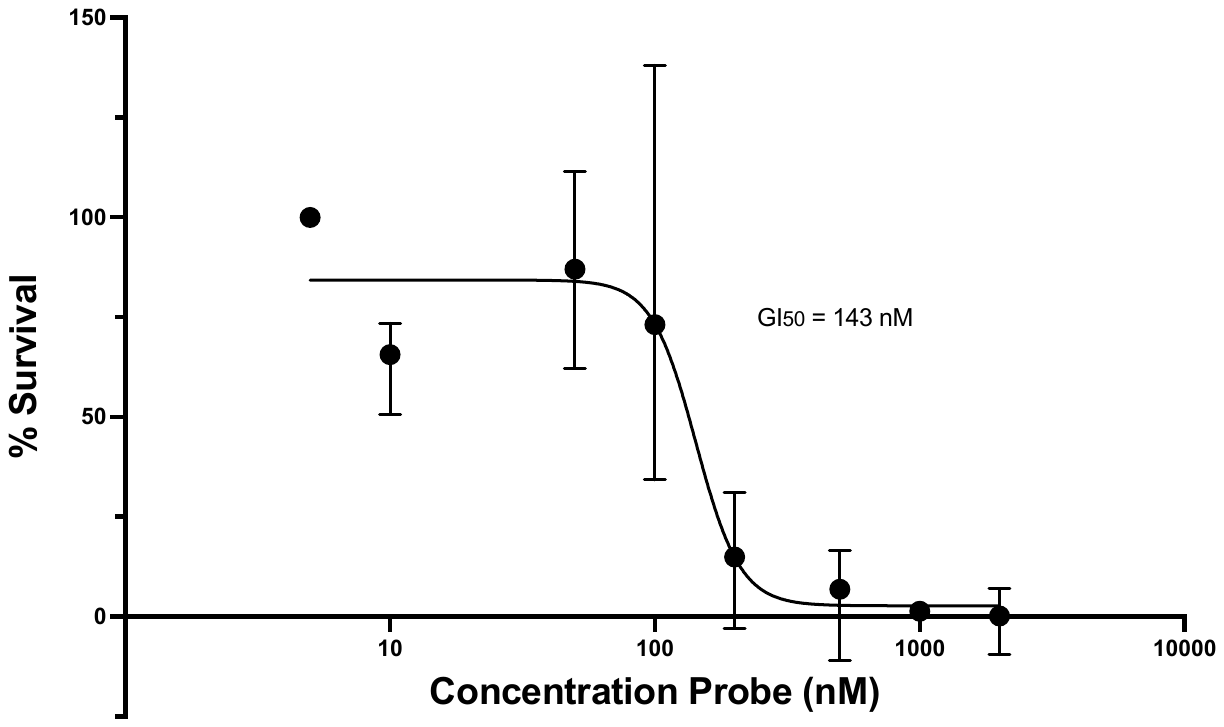

**Figure S10** – MTS assay of **TOR-G4** in U2OS cells measured (in triplicate) following 6-hour probe incubation; The resulting GI_50_ is 143 nM.

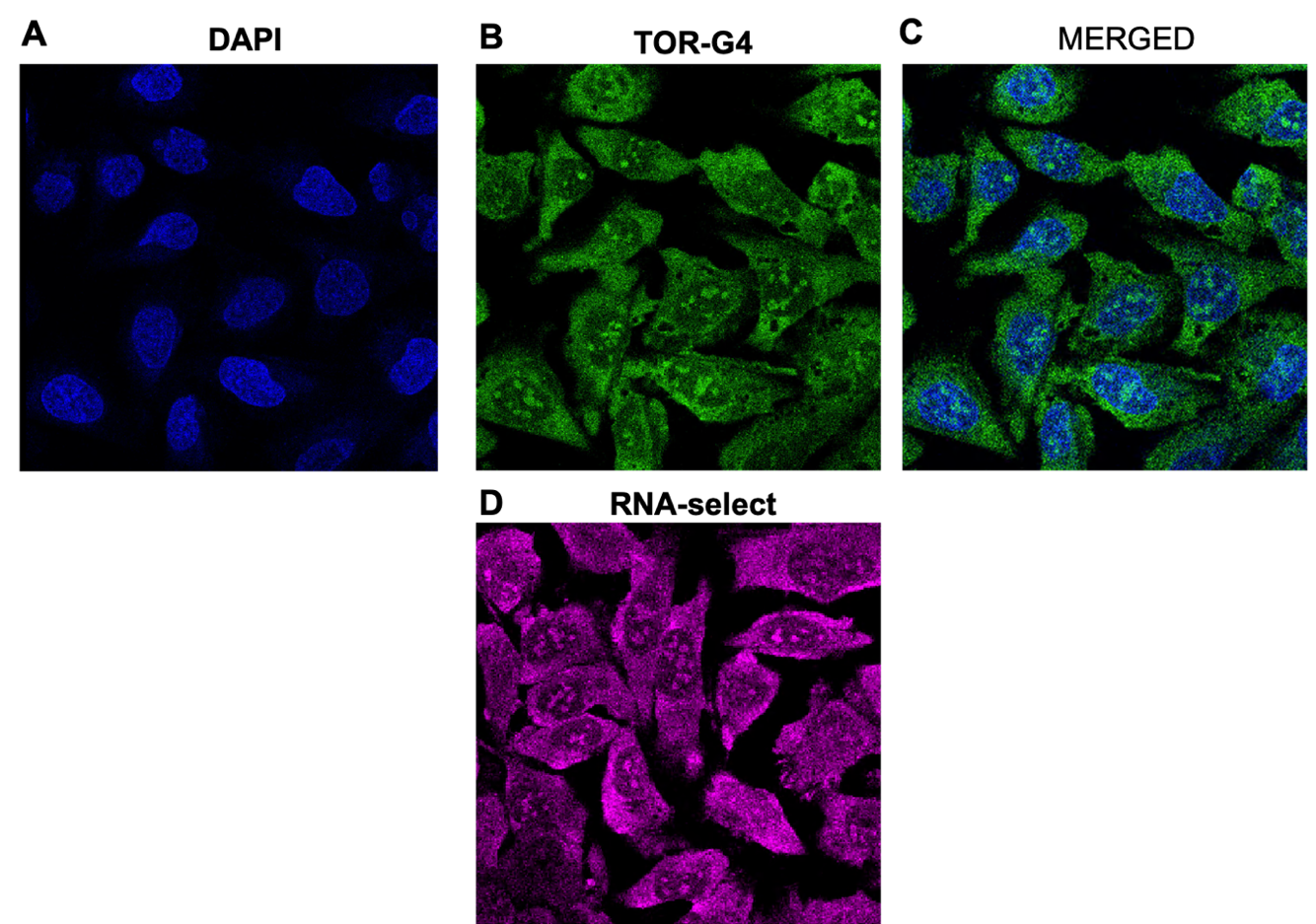

**Figure S11** – Staining of U2OS cells with A) DAPI (400-500 nm detection); B) **TOR-G4** (550- 700 nm detection) C) DAPI and **TOR-G4** merged D) SYTO RNASelect (550-700 nm detection).

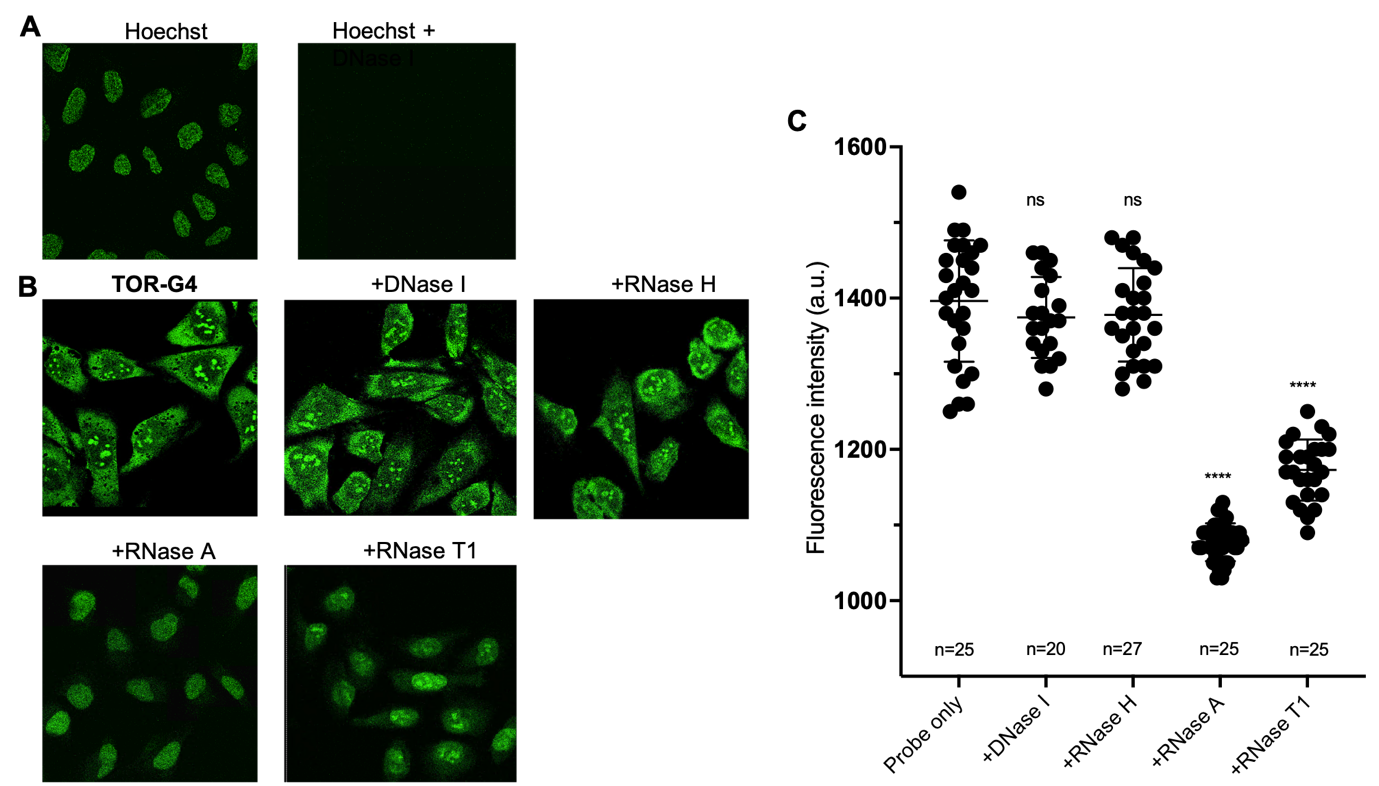

**Figure S12** – A) Confocal images of DNA stain Hoechst 33342 in U2OS before and after treatment with DNase I. Two-photon excitation at 760 nm, detection at 400-500 nm. B) Confocal images of **TOR-G4** in U2OS cells before and after treatment with nucleases. C) Quantification of fluorescence intensity of **TOR-G4** within U2OS cells before and after treatment with nucleases. Excitation at 477 nm, detection at 550- 700 nm. Statistical significance was assessed via Mann-Whitney u-test: ns = non-significant, **** = p<0.001.

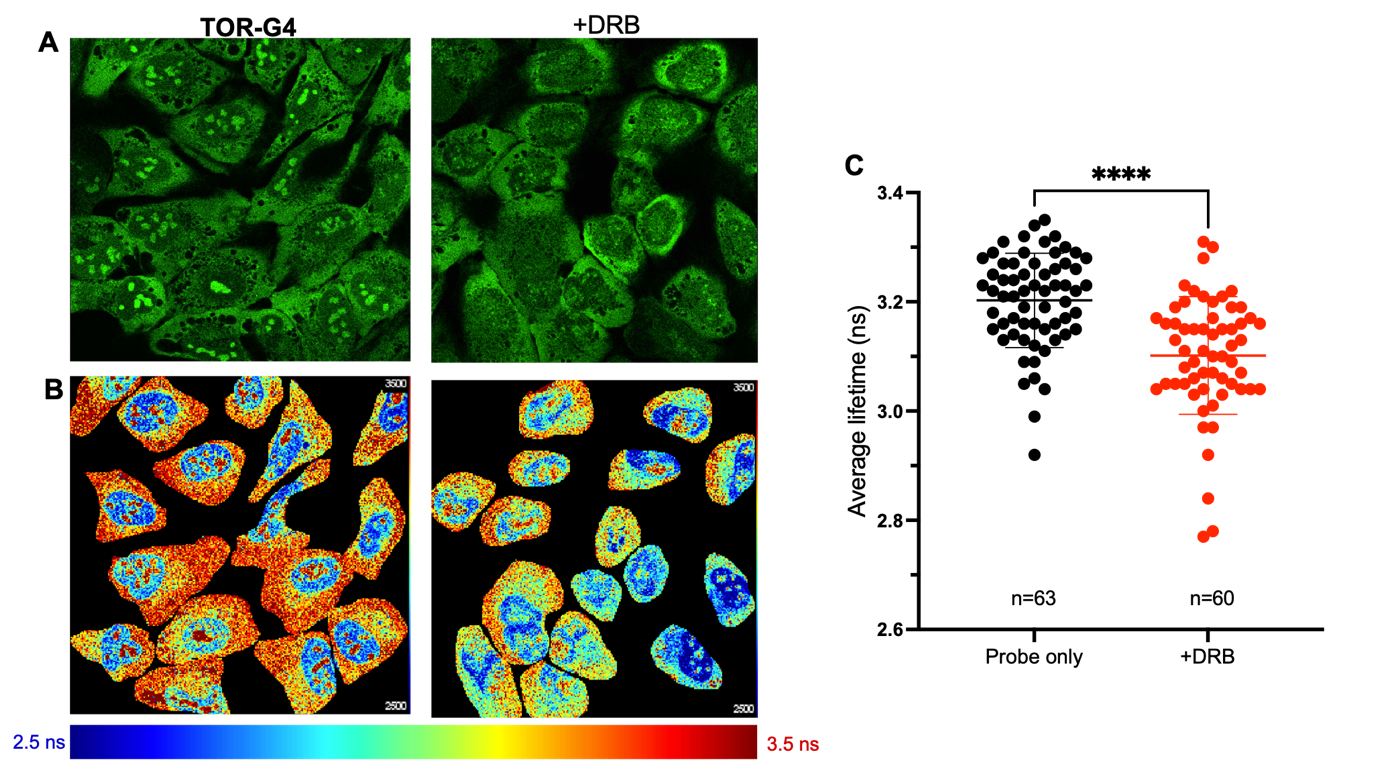
**Figure S13** – A) Confocal and B) FLIM images of **TOR-G4** in U2OS cells before and after transcriptional inhibition with DRB. C) Quantification of probe cellular lifetime before and after DRB treatment.

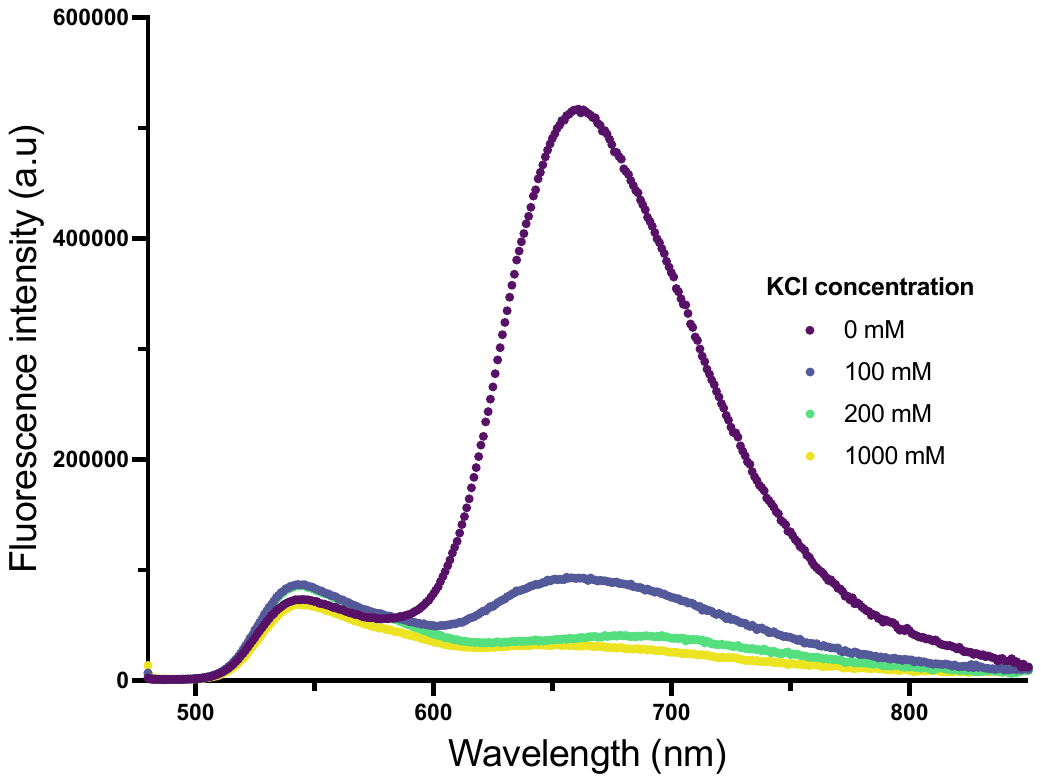

**Figure S14** – Emission spectra of **TOR-G4** (2 $\mu$M) recorded in aqueous buffered solution in the presence of c-MYC DNA (100 $\mu$g/mL) with increasing concentrations of KCl (0-1000 mM). The decrease in the intensity of the 660 nm band is consistent with disaggregation of **TOR-G4** upon increasing ionic strength, or K^+^ ions.

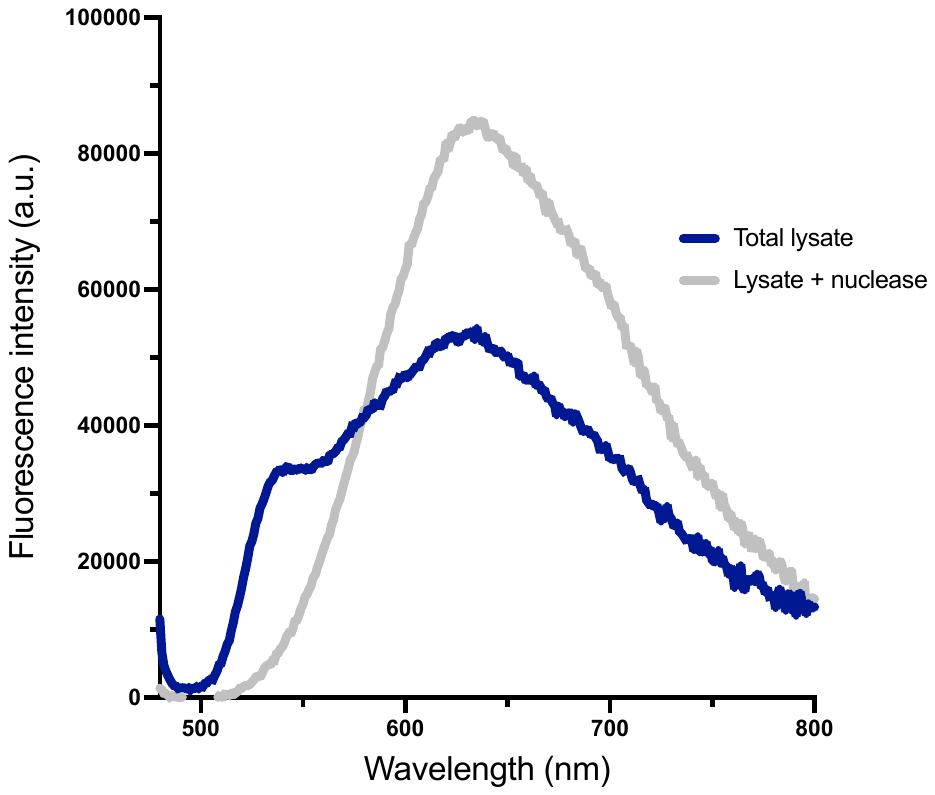

**Figure S15** – Emission spectra of **TOR-G4** (2 $\mu$M) in U2OS cell lysate before (blue) and after (grey) nuclease treatment. The emission of the monomeric form of **TOR-G4** at 540 nm disappears upon nuclease treatment, consistent with our conclusion that nucleic acids are required for the disaggregation of **TOR-G4**.

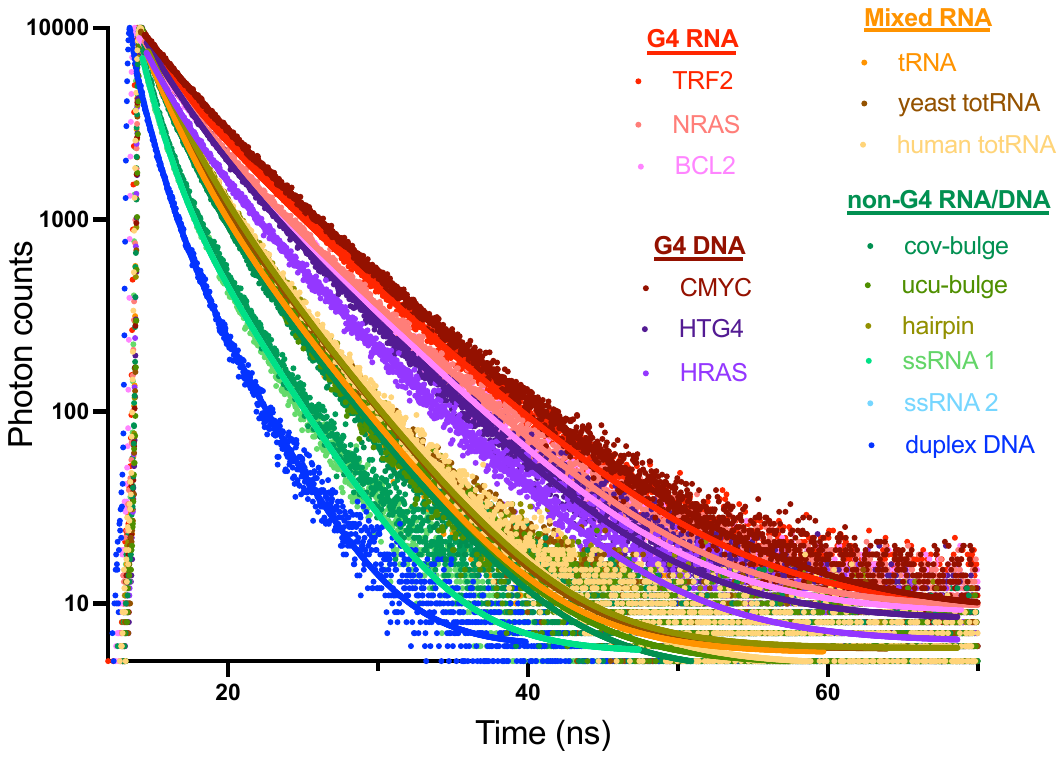

**Figure S16 –** Time-resolved fluorescence decay traces of **TOR-G4** (2 $\mu$M) bound to various DNA and RNA sequences (100 $\mu$g/mL). Excitation was at 470 nm, detection at 540 nm.

**Table S3** – Goodness of fit (chi-sq value) of **TOR-G4** decay to a biexponential function when bound to various DNA/RNA structures.

| Sequence | Chi-sq |
| --- | --- |
| CMYC | 1.15 |
| HRAS | 1.24 |
| HT-G4 | 1.20 |
| TRF2 | 1.16 |
| BCL2 | 1.14 |
| NRAS | 1.19 |
| human totRNA | 1.39 |
| yeast totRNA | 1.36 |
| tRNA | 1.23 |
| hairpin RNA | 1.30 |
| cov-bulge | 1.30 |
| ucu-bulge | 1.38 |
| ssRNA 1 | 1.42 |
| ssRNA 2 | 1.30 |
| duplex DNA | 1.31 |

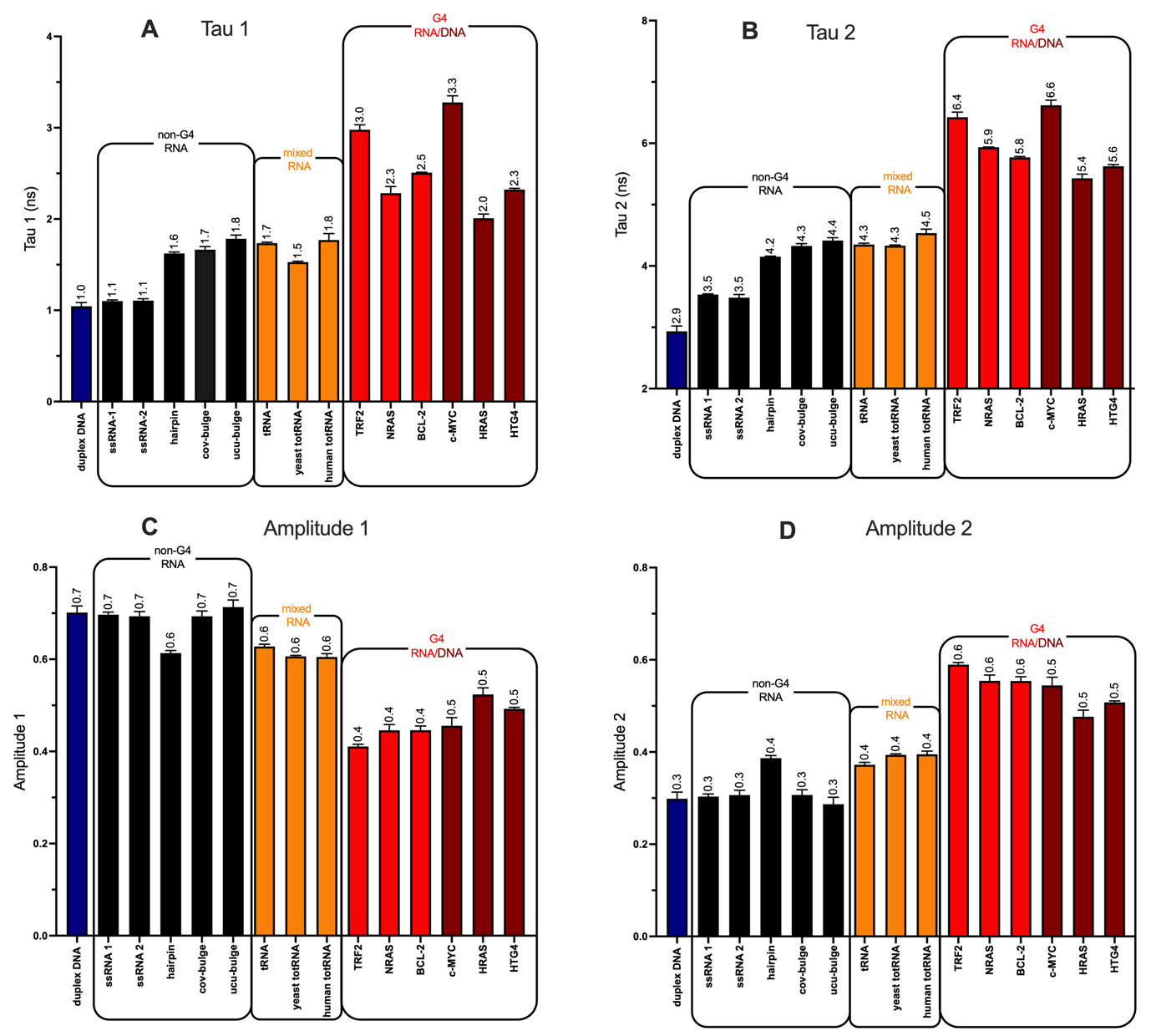

**Figure S17 –** Fluorescence lifetime component analysis of **TOR-G4** (2 $\mu$M) bound to various DNA and RNA topologies (100 $\mu$g/mL), showing lifetime components A) Tau1 and B) Tau 2, and decay amplitudes C) Amplitude 1 and D) Amplitude 2. Excitation at 470 nm, detection at 540 nm. The original time-resolved decays are shown in Figure S11.

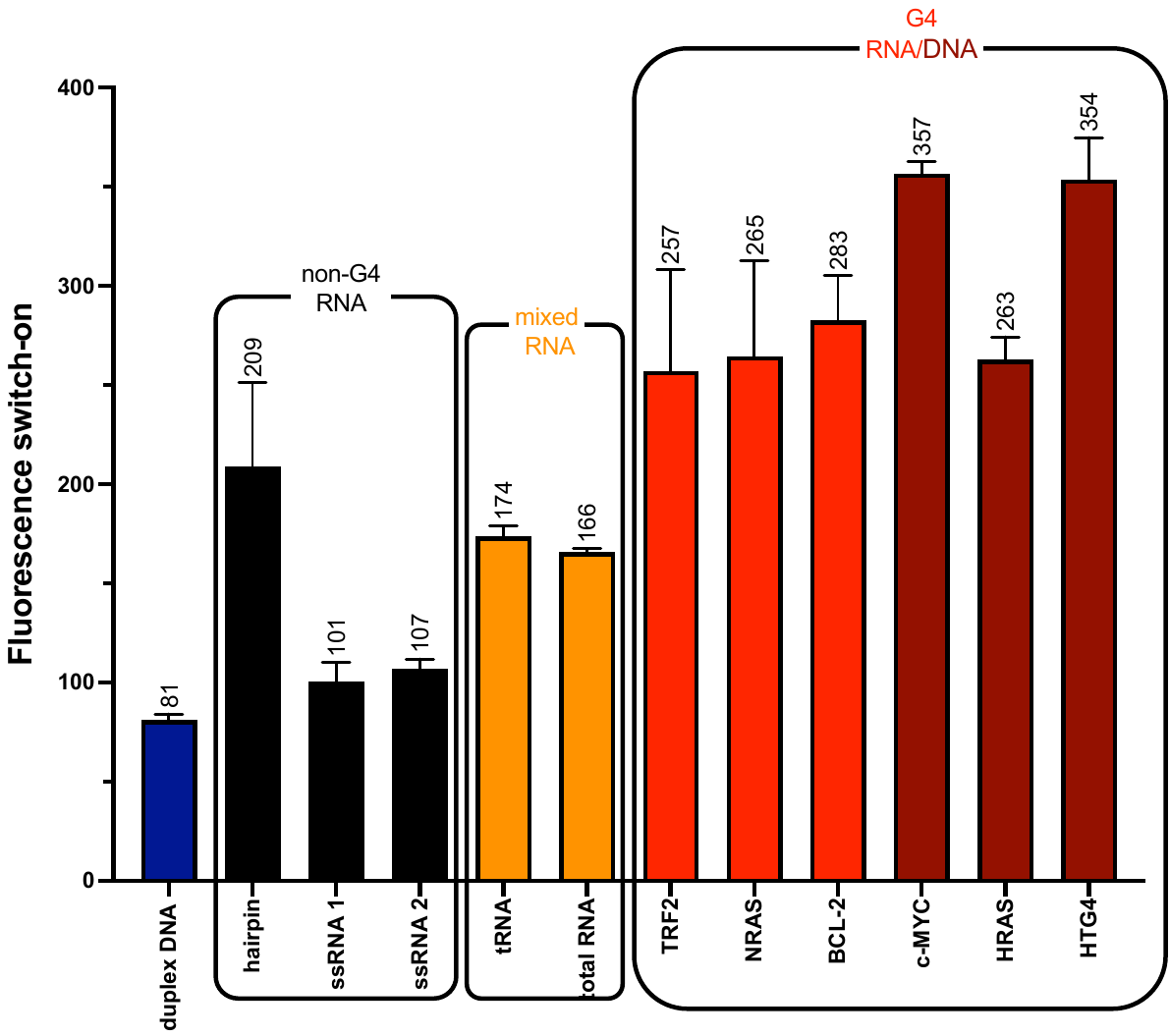

**Figure S18 –** Fluorescence switch-on (at 540 nm) of **TOR-G4** (2$\mu$M) when interacting with various DNA and RNA topologies (100 $\mu$g/mL)

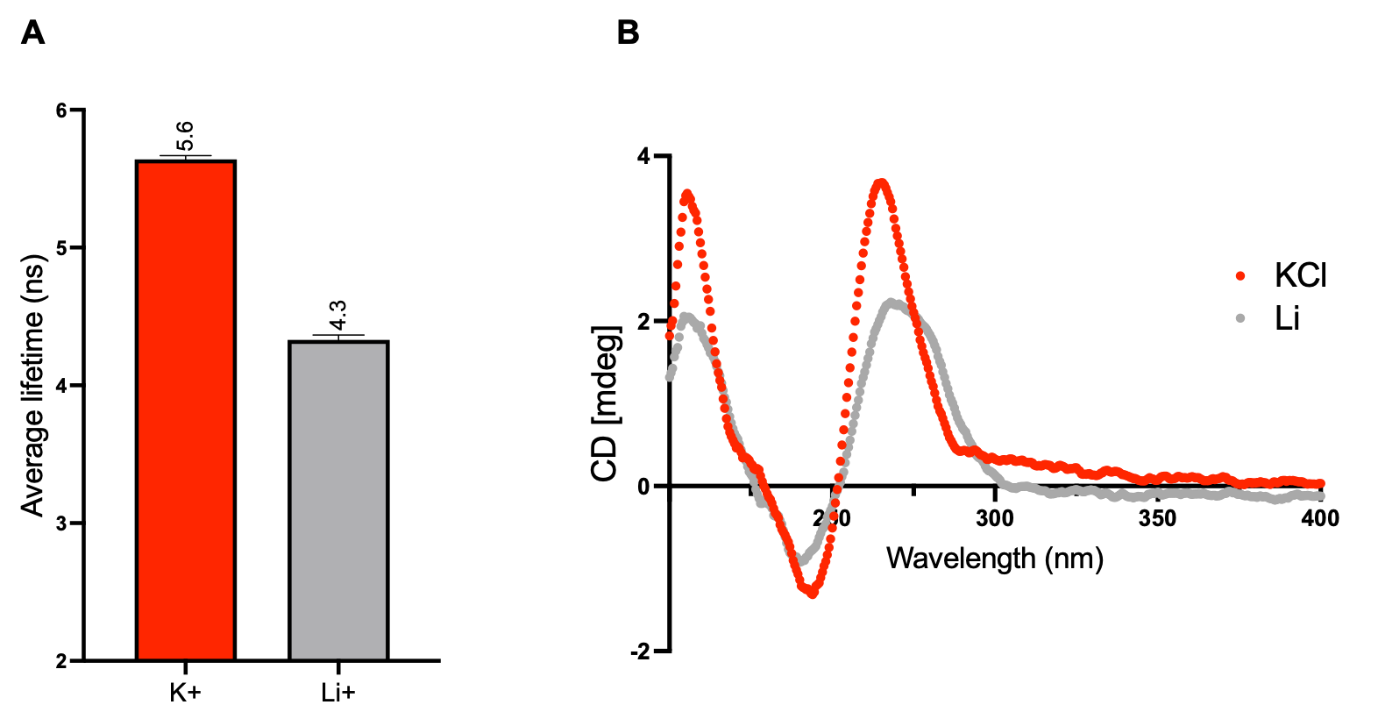

**Figure S19** – A) Average weighted lifetime of **TOR-G4** bound to *c-MYC* in K^+^- or Li^+^-containing aqueous buffer. Excitation at 470 nm and emission detected at 540 nm. B) CD spectra of *c-MYC* in K^+^- and Li^+^-containing aqueous buffer.

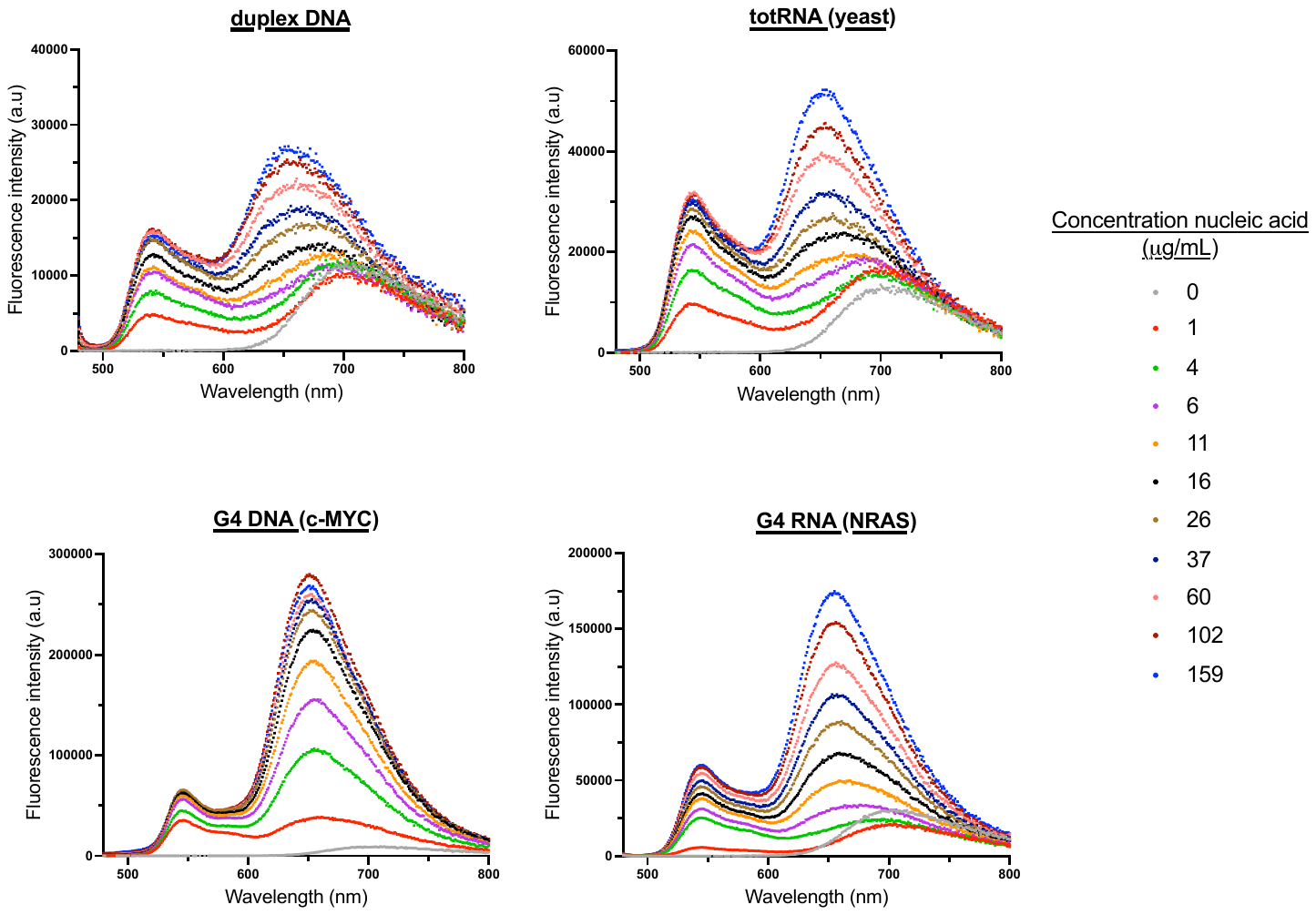

**Figure S20 –** Emission spectra of **TOR-G4** (2 $\mu$M) with increasing concentrations (0-159 $\mu$g/mL) of various DNA and RNA topologies. Excitation at 470 nm.

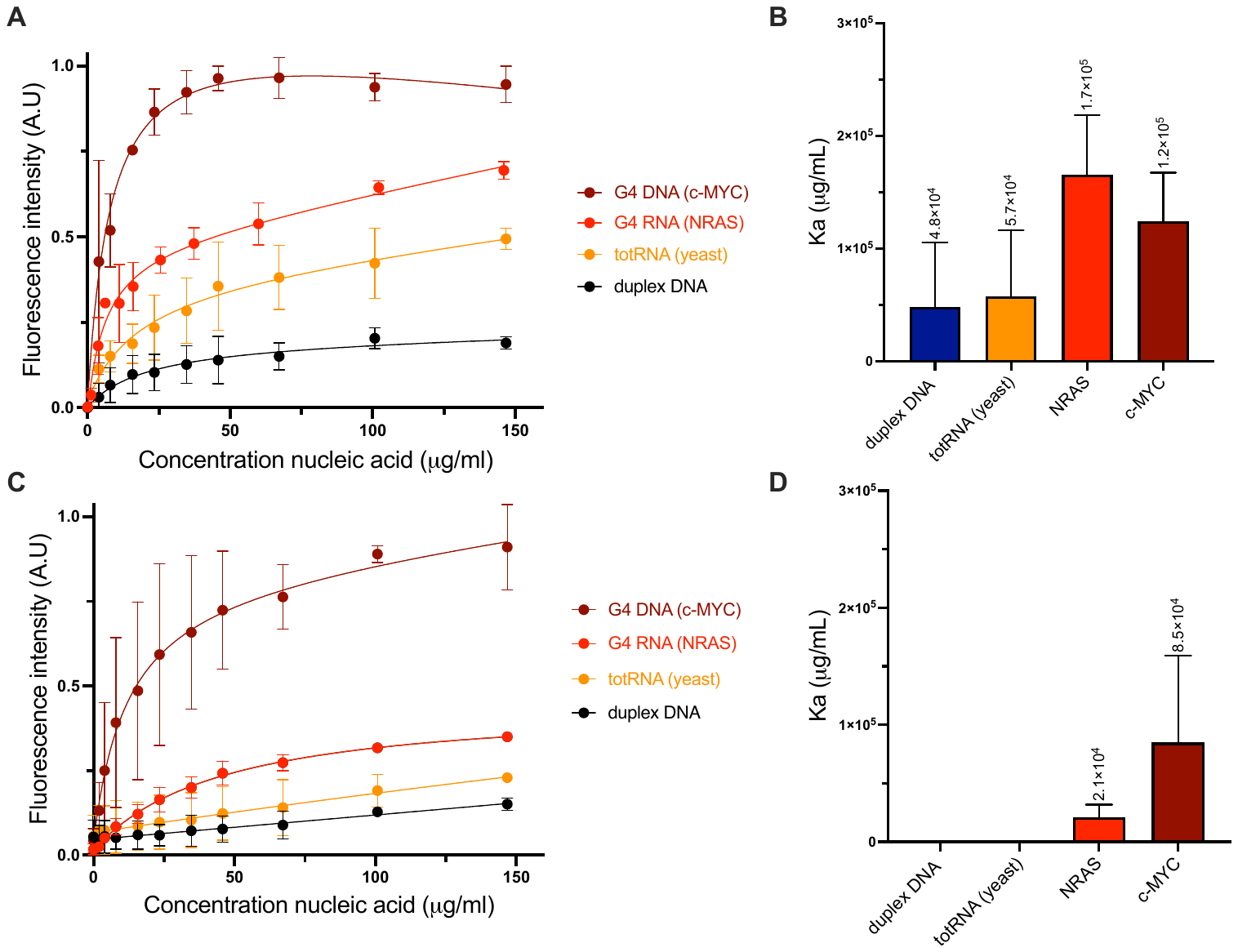

**Figure S21 –** A) Binding curves and B) apparent binding affinities of **TOR-G4** monomer (540 nm detection) obtained from binding titrations of **TOR-G4** (2 $\mu$M) with various DNA and RNA sequences. C) Binding curves and D) apparent binding affinities of **TOR-G4** dimer (660 nm detection) interacting with various nucleic acids. We note that it is not possible to fully deconvolute the effects of binding to nucleic acids from that of aggregation, which affect the absorbance and emission of the probe (see Figure S20). Error bars are standard deviation of experiments performed in triplicate. Associations constants calculated by fitting binding curves to Graphpad prism one site –total model.

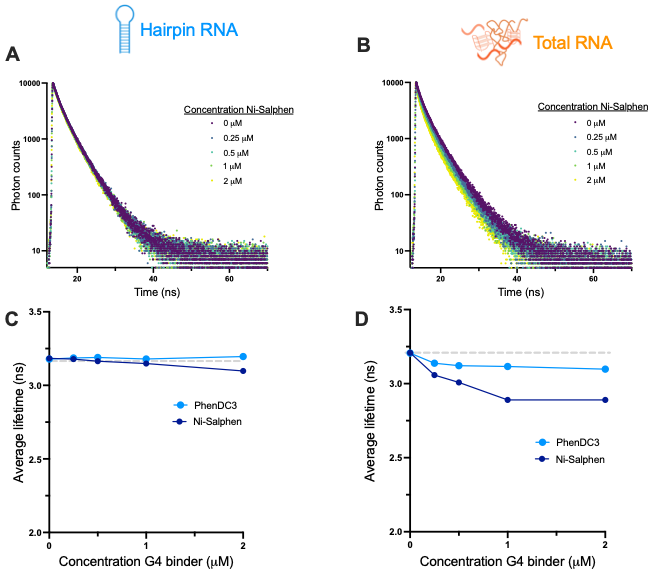

**Figure S22** – Time-resolved decays from displacement assay of **TOR-G4** (2 $\mu$M) from A) hairpin RNA and B) total RNA using G4 binder Ni-Salphen (0-2 $\mu$M). Average fluorescence lifetime of **TOR-G4** (2 $\mu$M) after displacement from C) hairpin RNA and D) total RNA using PhenDC3 and Ni-Salphen (0-2 $\mu$M).

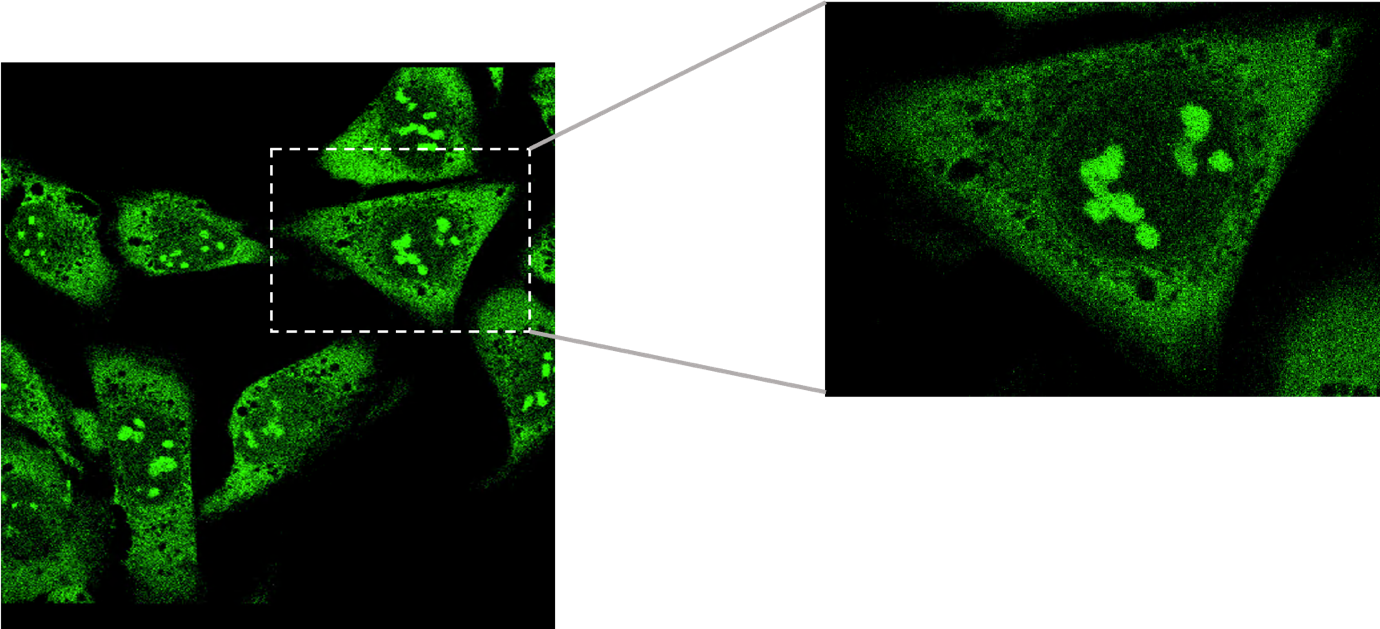

**Figure S23** – Confocal image of **TOR-G4** in U2OS cells, excitation at 514 nm and detection at 550-700 nm. Corresponding FLIM image in Figure 3A of main text.

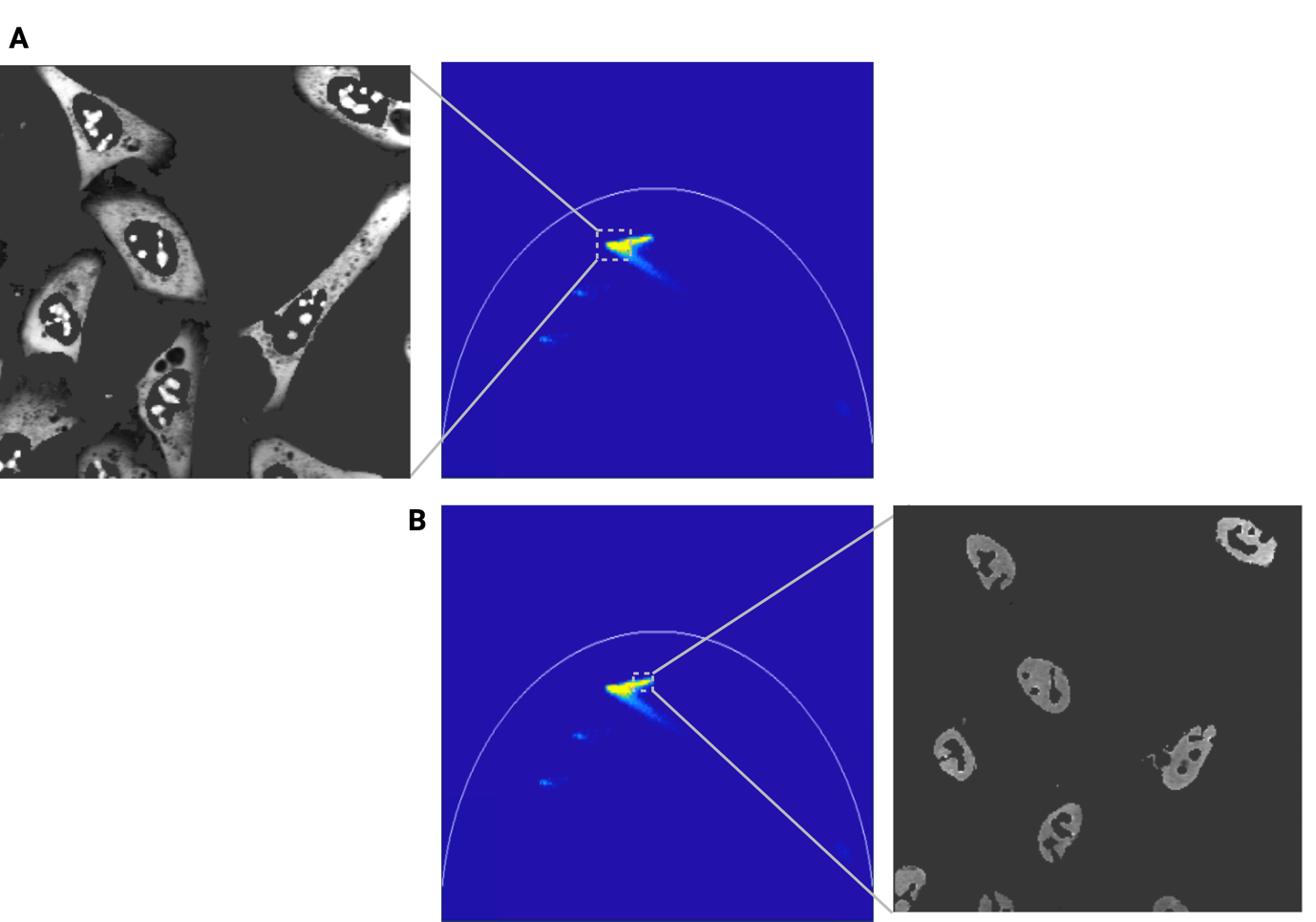

**Figure S24** - Phasor analysis of **TOR-G4** fluorescence decays within U2OS cells**.** A) Cellular segmentation of high lifetime portion of phasor plot. B) Cellular segmentation of low lifetime portion of phasor plot. Analysis performed in FLIMfit.

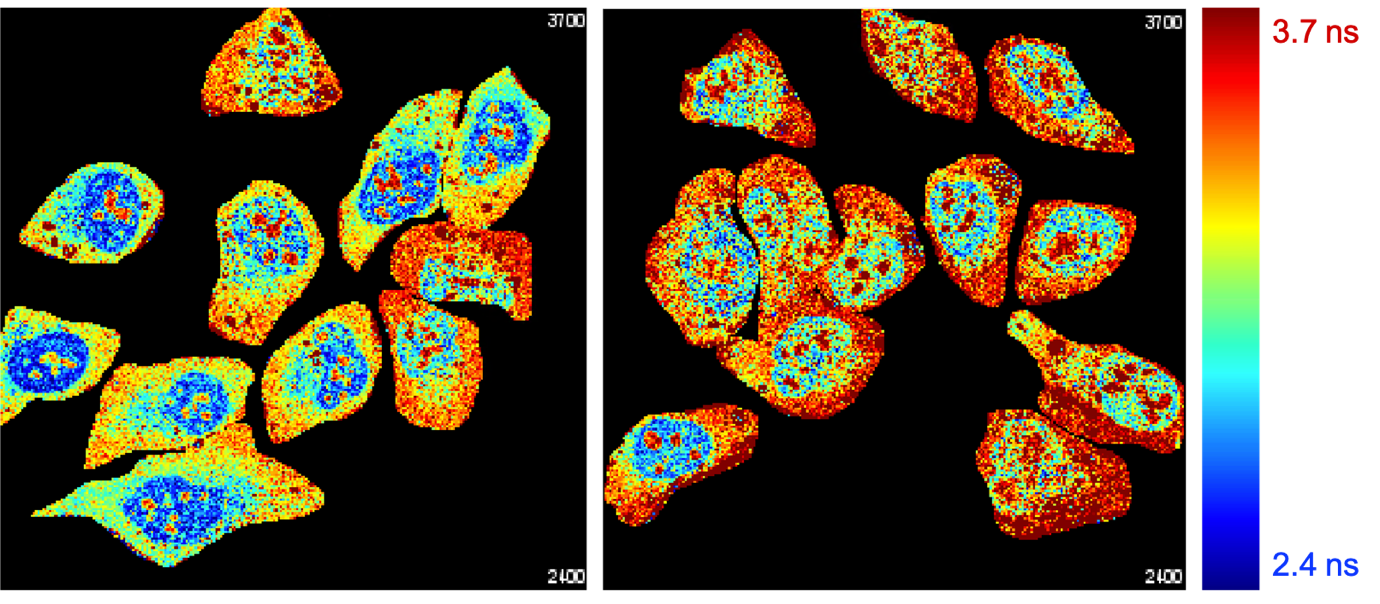

**Figure S25** – Example FLIM images of **TOR-G4** (5 $\mu$M) visualized in U2OS cells after two-photon excitation. Excitation at 760 nm, detection at 550 – 700 nm.

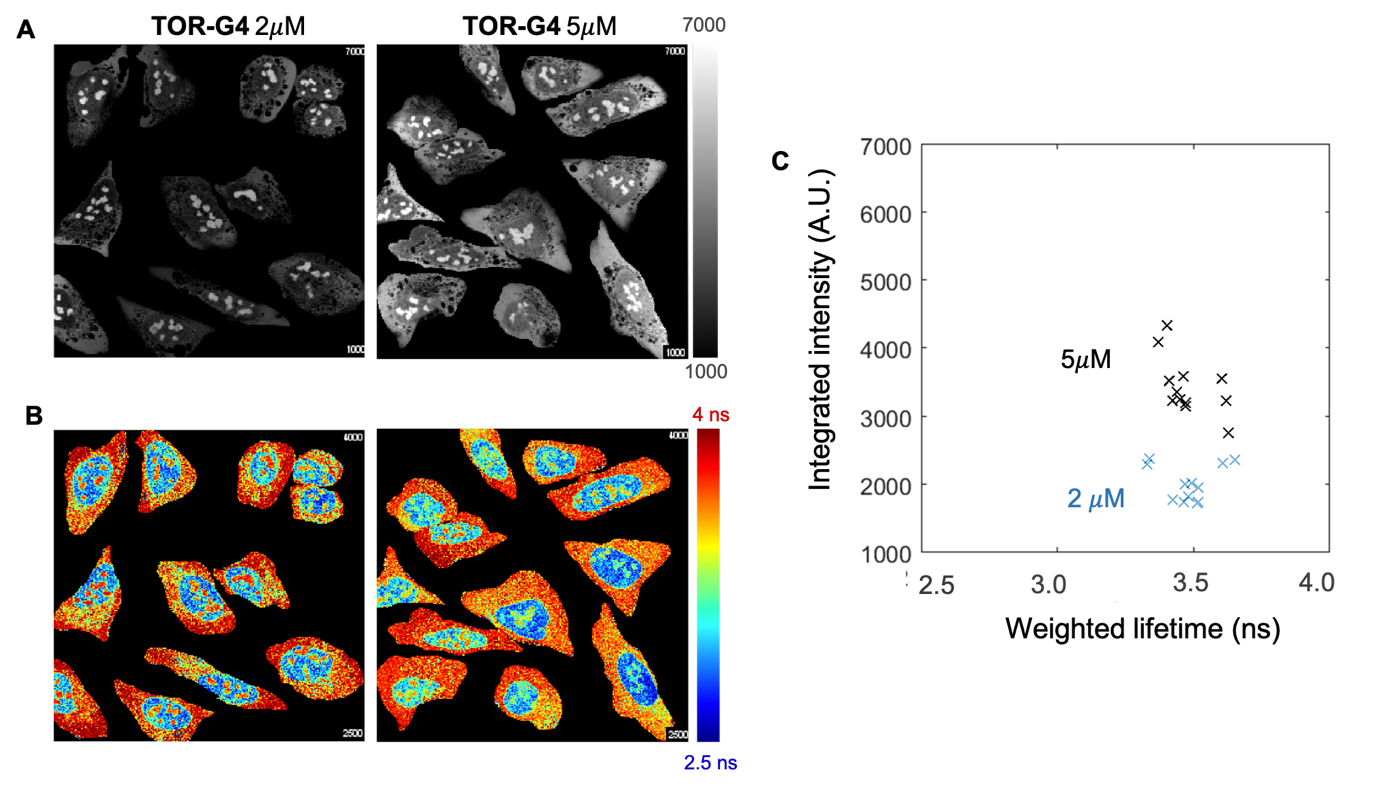

**Figure S26** – A) Confocal intensity and B) FLIM images of **TOR-G4** in U2OS cells incubated at 2 $\mu$M or 5 $\mu$M for 2 hours. C) Correlation between fluorescence intensity and fluorescence lifetime of cells incubated with either 2 $\mu$M (blue crosses) or 5 $\mu$M (black crosses) of **TOR-G4**. Excitation at 477 nm and detection at 550–700 nm.

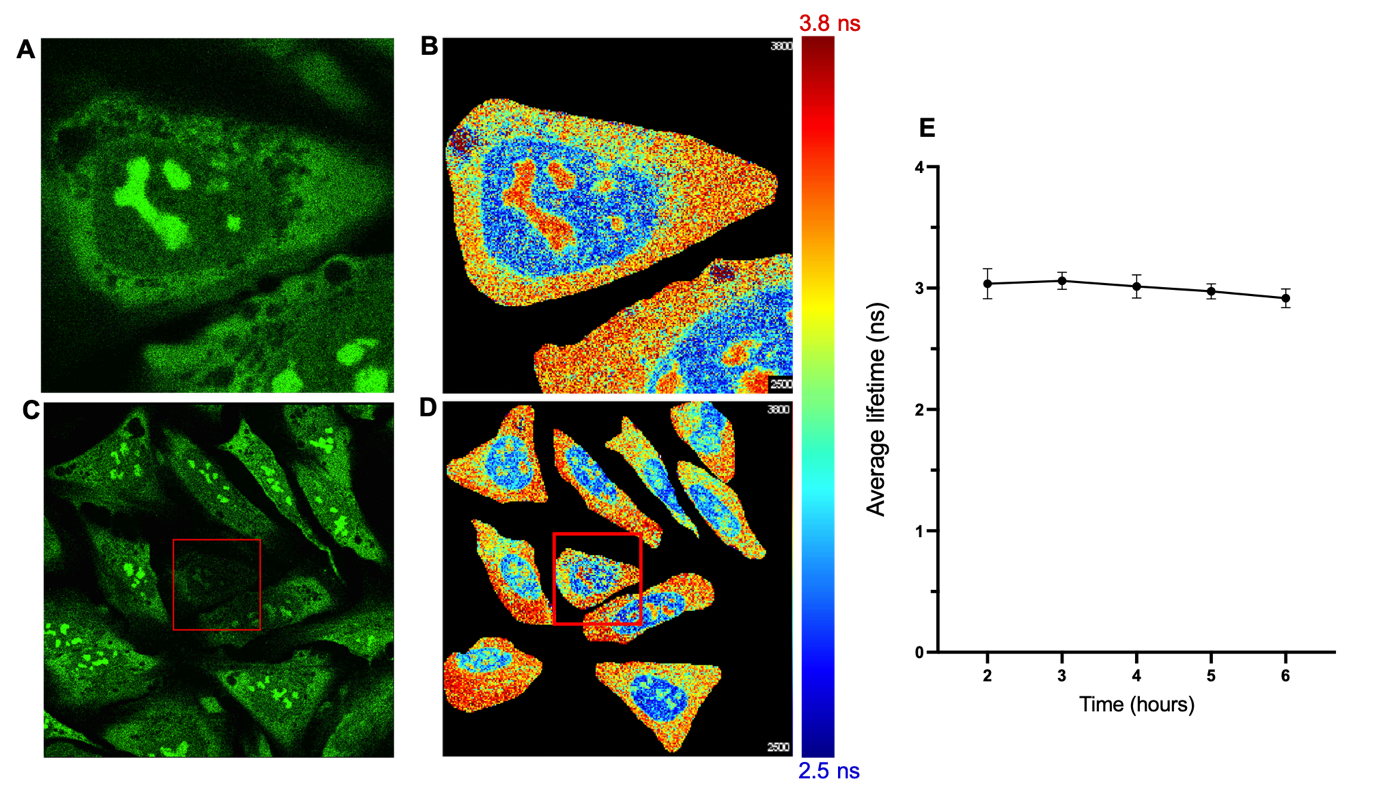

**Figure S27** – Investigating photostability of **TOR-G4.** A) Confocal image of cells prior to irradiation B) First FLIM image of irradiated cells (500s acquisition time). C) Confocal image of cells after first image acquisition – red box shows the irradiated area. D) Second FLIM image of cells – red box shows the previously irradiated area. E) Fluorescence lifetime of **TOR-G4** within U2OS cells across 6 hours of continuous imaging. Excitation at 477 nm and detection at 550–700 nm.

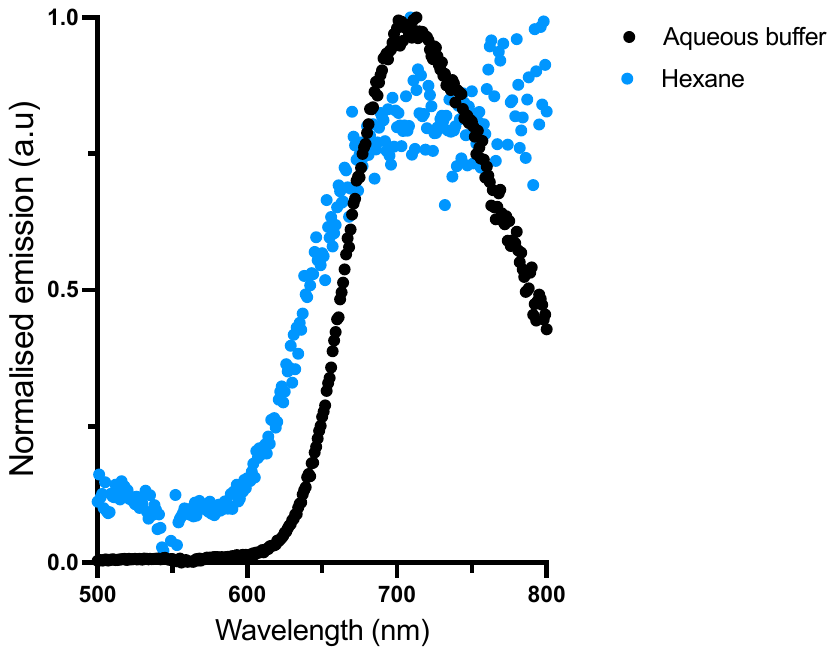

**Figure S28** - Emission spectra of **TOR-G4** in aqueous buffer (black) and hexane (blue). Excitation at 470 nm. No significant shift in the emission peak is observed when changing solvent, suggesting the emission is not influenced by charge transfer processes.

### 6. References

(1)	Lu, Y.-J.; Deng, Q.; Hou, J.-Q.; Hu, D.-P.; Wang, Z.-Y.; Zhang, K.; Luyt, L. G.; Wong, W.-L.; Chow, C.-F. Molecular Engineering of Thiazole Orange Dye: Change of Fluorescent Signaling from Universal to Specific upon Binding with Nucleic Acids in Bioassay. *ACS Chem. Biol.* **2016**, *11* (4), 1019–1029. https://doi.org/10.1021/acschembio.5b00987.

(2)	Abd Karim, N. H.; Mendoza, O.; Shivalingam, A.; Thompson, A. J.; Ghosh, S.; Kuimova, M. K.; Vilar, R. Salphen Metal Complexes as Tunable G-Quadruplex Binders and Optical Probes. *RSC Adv.* **2013**, *4* (7), 3355–3363. https://doi.org/10.1039/C3RA44793F.

(3)	Spielmann, H. P.; Wemmer, D. E.; Jacobsen, J. P. Solution Structure of a DNA Complex with the Fluorescent Bis-Intercalator TOTO Determined by NMR Spectroscopy. *Biochemistry* **1995**, *34* (27), 8542–8553. https://doi.org/10.1021/bi00027a004.

(4)	Dai, J.; Carver, M.; Hurley, L. H.; Yang, D. Solution Structure of a 2:1 Quindoline-c-MYC G-Quadruplex: Insights into G-Quadruplex-Interactive Small Molecule Drug Design. *J. Am. Chem. Soc.* **2011**, *133* (44), 17673–17680. https://doi.org/10.1021/ja205646q.

(5)	Trott, O.; Olson, A. J. AutoDock Vina: Improving the Speed and Accuracy of Docking with a New Scoring Function, Efficient Optimization, and Multithreading. *J. Comput. Chem.* **2009**, *31* (2), 455-461. https://doi.org/10.1002/jcc.21334.

(6)	Warren, S. C.; Margineanu, A.; Alibhai, D.; Kelly, D. J.; Talbot, C.; Alexandrov, Y.; Munro, I.; Katan, M.; Dunsby, C.; French, P. M. W. Rapid Global Fitting of Large Fluorescence Lifetime Imaging Microscopy Datasets. *PLoS One* **2013**, *8* (8), 70687. https://doi.org/10.1371/journal.pone.0070687.
